# Genetic determinants of COVID-19 drug efficacy revealed by genome-wide CRISPR screens

**DOI:** 10.1101/2020.10.26.356279

**Authors:** Wei Jiang, Ailing Yang, Jingchuan Ma, Dawei Lv, Mingxian Liu, Liang Xu, Chao Wang, Zhengjin He, Shuo Chen, Jie Zhao, Shishuang Chen, Qi Jiang, Yankai Chu, Lin Shan, Zhaocai Zhou, Yun Zhao, Gang Long, Hai Jiang

## Abstract

Immunomodulatory agents dexamethasone and colchicine, antiviral drugs remdesivir, favipiravir and ribavirin, as well as antimalarial drugs chloroquine phosphate and hydroxychloroquine are currently used in the combat against COVID-19^1–16^. However, whether some of these drugs have clinical efficacy for COVID-19 is under debate. Moreover, these drugs are applied in COVID-19 patients with little knowledge of genetic biomarkers, which will hurt patient outcome. To answer these questions, we designed a screen approach that could employ genome-wide sgRNA libraries to systematically uncover genes crucial for these drugs’ action. Here we present our findings, including genes crucial for the import, export, metabolic activation and inactivation of remdesivir, as well as genes that regulate colchicine and dexamethasone’s immunosuppressive effects. Our findings provide preliminary information for developing urgently needed genetic biomarkers for these drugs. Such biomarkers will help better interpret COVID-19 clinical trial data and point to how to stratify COVID-19 patients for proper treatment with these drugs.

## Introduction

In response to the surging COVID-19 pandemic, several drugs have been reported to show efficacy in COVID-19 patients, but clear conclusions on some of them are still lacking. Among these drugs, dexamethasone and colchicine serve to suppress excessive immune response that damages tissues and organs in severe COVID-19 cases. Antiviral drugs such as remdesivir, favipiravir and ribavirin interfere with viral RNA polymerase and inhibit the replication of SARS-CoV-2. Antimalarial drugs chloroquine phosphate and hydroxychloroquine, whose use in COVID-19 is more controversial, suppress both immune response and viral replication. Currently little is known about the genetic biomarkers that modulate these drugs’ efficacy. Such knowledge is urgently needed to better manage the COVID-19 pandemic.

Several groups of cellular proteins may affect drug efficacy. The list includes drug transporters, drug metabolizing enzymes, proteins that compete with the intended therapeutic target for drug binding, as well as proteins that mitigate or exacerbate drug-induced damage. Such proteins may differentially regulate the cellular stresses caused by a drug. If the stresses caused by COVID-19 drugs could result in massive cell death, it will be possible to capture genes that modify drug efficacy by analyzing cells that survived drug treatment (Fig. 1a).

**Figure 1.**
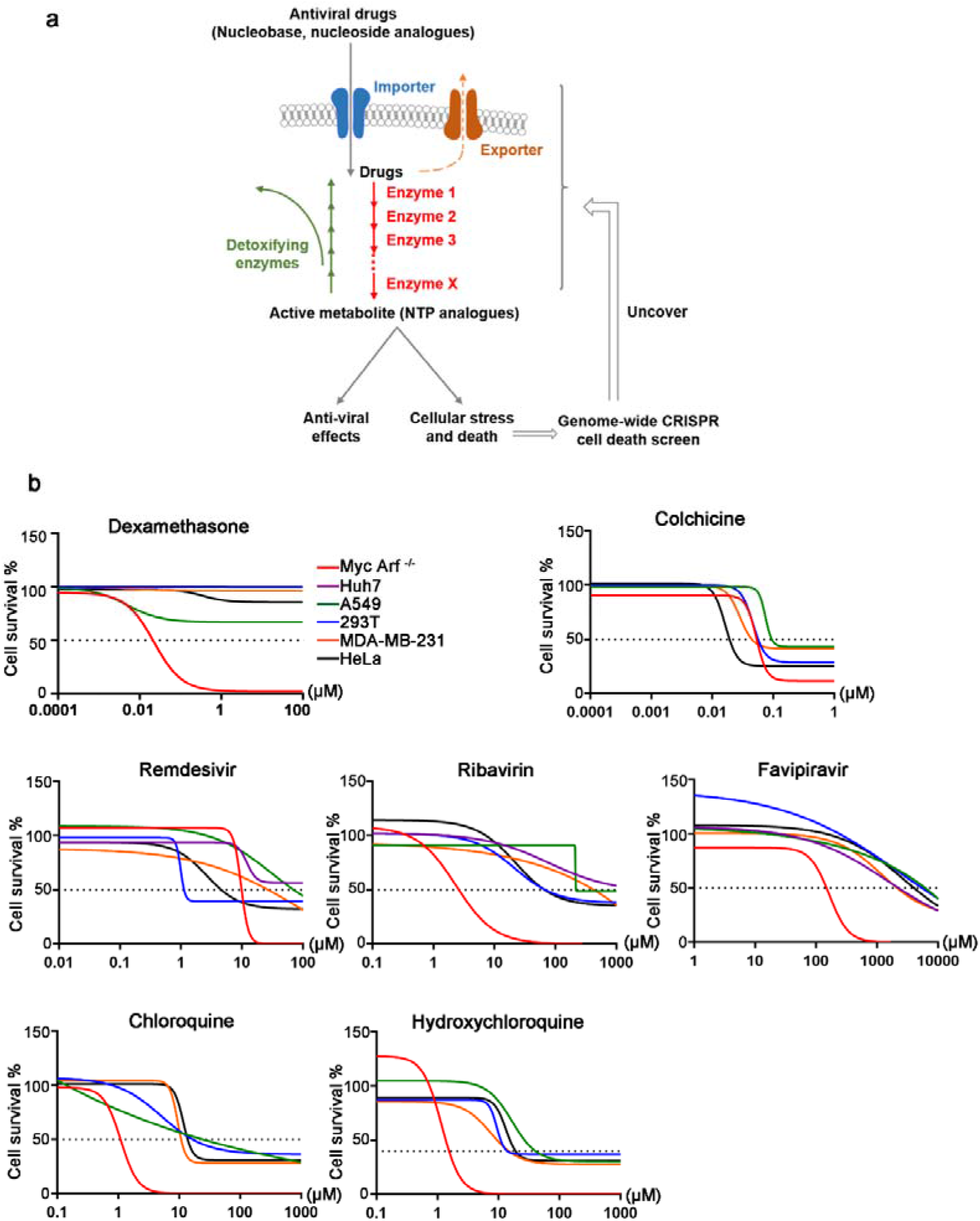
A genome-wide screen approach to identifying genes that modify the efficacy of COIVD-19 drugs. **a)** A diagram depicting the rationale of the screen. For example, antiviral drugs suppress viral replication by generating NTP analogs, which inhibit viral RNA polymerase. Such NTP analogs also interfere with cellular process that utilize normal NTPs, causing cellular stress that are sufficient to induce death in Eμ-Myc;Arf−/− cells. By introducing genome-wide sgRNA library into such cells and analyzing which gene knockout enabled survival after antiviral drug treatment, it is possible to uncover genes that mediate the transport of these drugs, as well as genes crucial for generating NTP analogs. Both classes of genes will also modify these drugs’ antiviral efficacy. **b**) COVID-19 drugs efficiently killed Eμ-Myc;Arf−/− cells, but not other commonly used cell lines.

However, most of the drugs used in COVID-19 treatment are not cytotoxic. They cannot efficiently kill commonly used cell lines even at very high concentrations (Fig. 1b), making it very difficult to conduct genome-wide screens. For antiviral drugs, there are additional difficulties that prevented large-scale screens. For example, if a sgRNA-mediated gene knockout in a host cell makes remdesivir more effective at suppressing viral replication, there is just no effective means to isolate that cell from other cells. Therefore, looking at drug’s direct impact on viral replication *per se* in a pool-based large screen is hardly executable. As a result, no genome-wide screen studies on antiviral drugs have been reported and there is very little knowledge on genetic biomarkers for these groups of important drugs. In this report, we developed a screen platform that could effectively solve the above problems. Using such a platform, we performed genome-wide screening on COVID-19 drugs and identified genes that critically regulate these drugs’ activities. Screen results for these drugs are presented separately, with validation experiments shown at the end of this report.

## Results

### Screen system

The Eμ-Myc;Arf−/− cell line represents a good model system for interrogating gene-drug interactions^17–21^ and has led to renewed understanding of several important drugs^19,21^. Importantly, with regard to drug screen, these cells are highly apoptosis-prone and readily undergo cell death at stress levels that cannot kill most other types of cells (Fig. 1b). The ability of Eμ-Myc;Arf−/− cells to amplify drug-induced perturbance into a cell death response is essential for the proposed screen (Fig. 1a), as it permits the extension of genome-wide screens onto many types of nontoxic drugs, including seven COVID-19 drugs in this study.

Using a retroviral backbone, we introduced a genome-wide sgRNA library ^22^ into cas9-expressing Eμ-Myc;Arf−/− cells at 200-fold coverage. For the vast majority of genes, there were 10 different sgRNAs per gene in the screen. Around 25 million of sgRNA-expressing cells were subjected to two rounds of treatment with a COVID-19 drug. Drug doses were adjusted so that approximately 10% to 20% of cells were alive at the lowest viability point (Extended Data Fig. 1). By examining which sgRNAs were highly enriched or depleted in surviving cells, we identified genes that modify the efficacy of these COVID-19 drugs.

### Colchicine

Colchicine is an immunomodulatory agent used to suppress inflammation in gout, and more recently shown to greatly reduce cardiovascular events and death in chronic coronary disease^23^. Multiple studies showed colchicine significantly lowered death rate in COVID-19 patients ^1–3^. Mechanistically, colchicine destabilizes microtubule and damages mitotic spindles, thereby ceasing the proliferation of immune cells and alleviating COVID-19 symptoms. In our genome-wide screen, we identified a number of genes whose sgRNAs were significantly enriched after colchicine treatment (Fig. 2a). Showing the validity of this screen system, ranked at No.10 was KIF2C (Fig. 2a’), a microtubule depolymerizing protein previously known to regulate the toxicity of microtubule destabilizers^24^. The top hit gene in our screen was TMEM161B, an uncharacterized protein localized primarily at the plasma membrane. In addition, both FLCN and FNIP were identified as top hits in our screen, suggesting the FLCN-FNIP protein complex may regulate colchicine-induced cell death. Other top hits include RQCD1, PTBP1, KMT2D, AP1G1, POU2F1 and BRD7 (Fig. 2a’), most of which have not been previously implicated for microtubule drugs.

**Figure 2.**
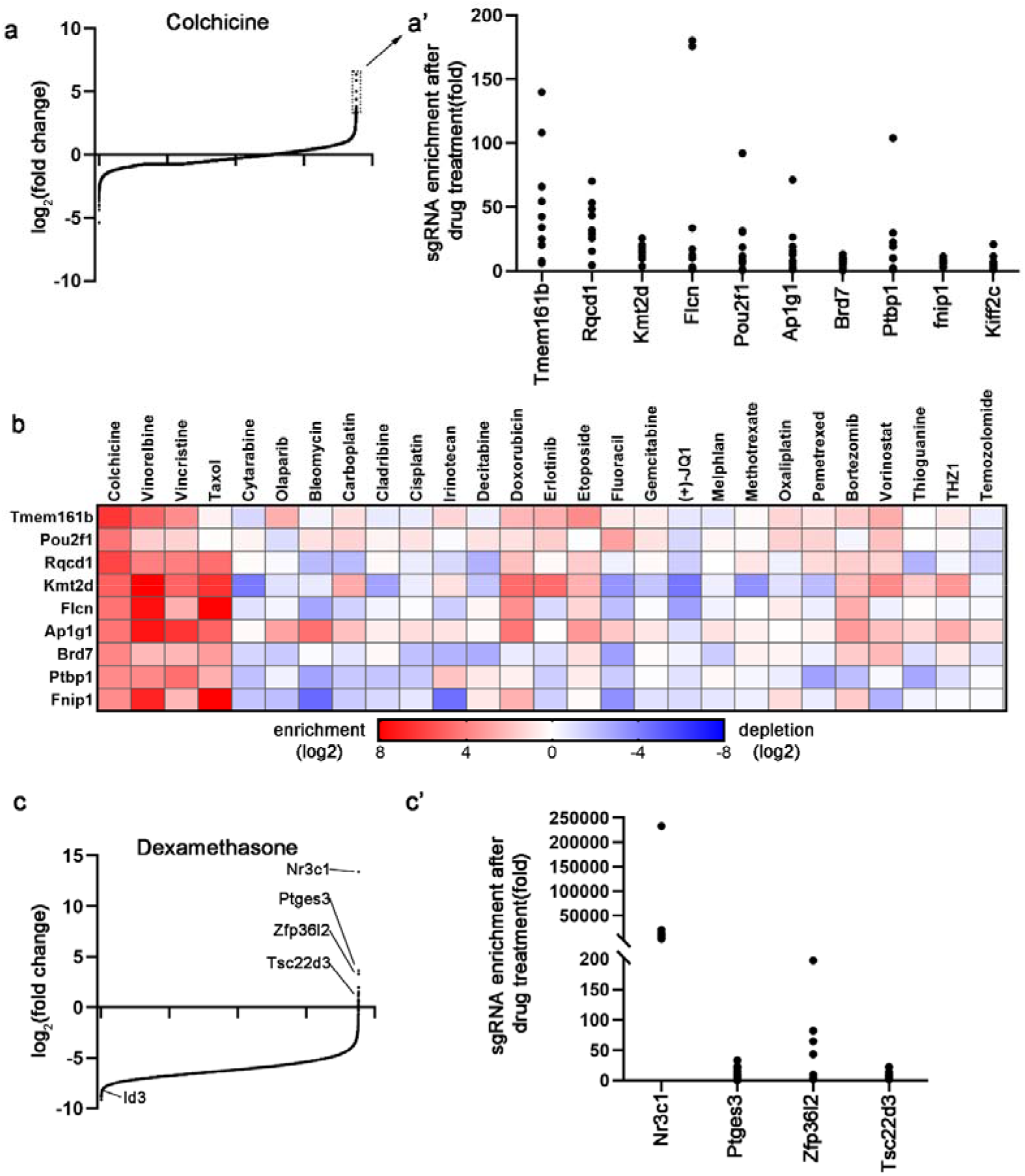
Genes enriched in colchicine and dexamethasone screens. **a and c)** Figures showing the overall distribution patterns of genes in the drug screen. Genes are sorted based on the fold change of sgRNAs after drug selection. Those most enriched genes are located at the far right. Top enriched genes are boxed out and detailed in **a’ and c’)**, which show the extent of enrichment of individual sgRNAs after drug selection. In **A’ and C’)**, each dot represents a sgRNA targeting the indicated gene. p values for all genes depicted in **a’ and c’)** are lower than 2.6×10^−7^. In **c)**, p value for Id3 depletion is 0.0007. **b)** How genes enriched in colchicine screen behaved in other drug screens conducted in our lab.

Prior to this study, we have completed genome-wide screens on taxol, vincristine and vinorelbine, three anticancer drugs that also target microtubule. To understand the performance of this screen system and whether certain genes are specific to colchicine, we referenced the other three screen results (Fig. 2b). We found that TMEM161B and POU2F1 affect cell viability in response to microtubule destabilizers colchicine, vincristine and vinorelbine, but not to taxol, which is a microtubule stabilizing drug. Other colchicine screen hit genes yielded similar phenotypes in all four microtubule drug screens, suggesting they are common crucial determinants of cell fate in response to microtubule disruption. Of note, these genes did not show up in our screens of other types of anticancer drugs (Fig. 2b). This analysis showed that for drugs of similar mechanisms of action, this screen system can consistently identify the same group of genes, highlighting Eμ-Myc;Arf−/− cell as a high performance screen platform. Importantly, the genes identified in our screen for the first time pointed to a group of potential biomarkers for colchicine’s use. If a COVID-19 patient carries genetic deficiency on one of these genes, colchicine may not sufficiently suppress the immune response to alleviate symptoms. This screen also shed light on the cytotoxic mechanisms of microtubule poisons, an important group of anticancer drugs whose death mechanisms are very unclear.

### Dexamethasone

Dexamethasone is a glucocorticoid that suppresses lymphocyte proliferation. It is used to treat many types of autoimmune diseases and hematological malignancies. Multiple studies demonstrated its efficacy in lowering death rate of COVID-19 patients^4–6^. As a cell line of B lymphocyte lineage, Eμ-Myc;Arf−/− cell serves as an ideal platform for interrogating dexamethasone’s immunosuppressive mechanism. In dexamethasone screen, sgRNAs targeting transcription factor NR3C1, the glucocorticoid receptor for dexamethasone, were dramatically enriched (Fig. 2c). Such a finding is consistent with previous knowledge that dexamethasone binding of NR3C1 initiates a transcription program that is highly detrimental to lymphocytes^25^. Our screen also recovered PTGES3, a NR3C1 chaperone protein required for glucocorticoid response^26^, and TSC22D3, which is an immunosuppressive gene induced by glucocorticoids^27^ (Fig. 2c). These results confirmed that the NR3C1-centered transcriptional program is central to dexamethasone’s immune suppressive effect. Considering this, we further asked whether certain cellular proteins with transcriptional inhibitory activities might antagonize such a transcription program and protect lymphocytes from dexamethasone-induced cell death. sgRNAs against such genes will be significantly depleted after dexamethasone selection. We identified one gene that fits such category. In the dexamethasone screen, inhibitor of DNA Binding 3 (ID3) was one of the most severely depleted genes. ID3 is capable of inhibiting transcriptional activity and is important for the B cell lineage. It is possible that ID3 may serve to limit the dexamethasone-NR3C1 transcription response, thereby protecting B cells from dexamethasone. Taken together, NR3C1, PTGES3, TSC22D3 and ID3 may represent candidate genetic biomarkers for dexamethasone’s usage in various diseases including COVID-19.

### Ribavirin

Ribavirin is a nucleoside analog that exhibits broad-spectrum antiviral activities. A recent study showed that a triple-drug combination of ribavirin, lopinavir-ritonavir and interferon β1, or the ribavirin lopinavir-ritonavir combination significantly accelerated viral clearance compared with lopinavir-ritonavir alone^7^. This suggests ribavirin may exert anti-SARS-CoV-2 activity in patients.

To suppress virus replication, ribavirin needs to be gradually converted to NTP analog to gain the ability to inhibit RNA-dependent RNA Polymerase (RdRp). Importantly, the same antiviral NTP analog will also interfere with vital cellular proteins including eIF4E and IMPDH^28^ and become detrimental to mammalian cells. Consequently, a cell death-based screen, enabled by Eμ-Myc;Arf−/− cells (Fig. 1b), will help identity enzymes that are important for the NTP conversion of ribavirin, which will also shed light on the antiviral effects of ribavirin (Fig 1a). Indeed, the top hit in our ribavirin screen was adenosine kinase (Adk) (Fig. 3a), an enzyme that catalyzes the formation of ADP from adenosine. This is consistent with a previous report that ADK is a key determinant for the anti-HCV activity of ribavirin^29^. As a proof of principle result, such an observation shows the validity of our screen rationale. Our screen uncovered two additional enzymes involved in nucleotide metabolism (Fig. 3a). Adenylosuccinate Synthase (ADSS) catalyzes the first step of the purine salvage pathway to form AMP. NUDT2 is a phosphatase that hydrolyzes dinucleotides to generate ATP or GTP. sgRNAs against these two genes were significantly enriched in cells that survived ribavirin treatment. This suggests that similar to ADK, both ADSS and NUDT2 may be critical to the NTP conversion and antiviral activity of ribavirin in cells. It may be interesting to analyze the SNP, copy number and expression level of these genes in responders and non-responders of ribavirin to explore potential biomarkers for COVID-19 ribavirin therapy.

**Figure 3.**
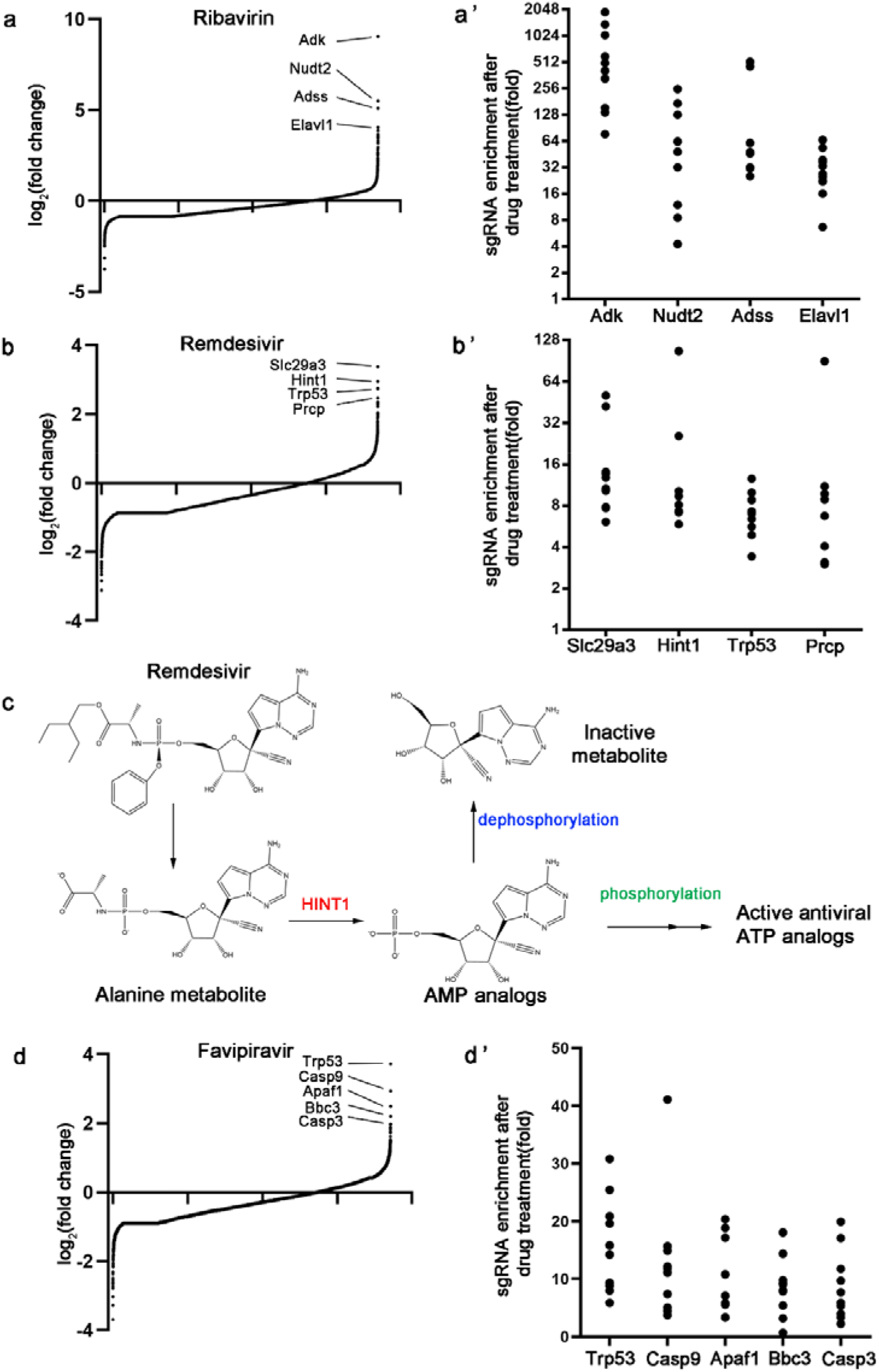
Genes enriched in antiviral drug screens. **a, b and d)** Figures showing the overall distribution patterns of genes in the drug screen. **a’, b’ and d’)** Figures showing the extent of enrichment of individual sgRNAs after drug selection. p values for all genes depicted in these panels are lower than 2.6×10^−7^. **c).** A diagram showing the metabolic process of remdesivir inside cells. One of the activation steps is carried out by HINT1.

### Remdesivir

Currently, remdesivir is the only antiviral drug approved by FDA for COVID-19 treatment. Earlier *in vitro* studies and subsequent compassionate use of remdesivir indicated its potential efficacy in treating COVID-19^30–32^. A randomized clinical trial in patients with severe disease failed to show superior outcome of remdesivir over control treatment^8^, whereas other studies suggested remdesivir could speed up recovery in COVID-19 patients ^9,10^. In the latter studies, remdesivir closely missed the significance threshold in terms of reduction of death rate (7.1 % with remdesivir vs 11.9% with placebo). It is therefore interesting to explore whether certain genetic backgrounds render remdesivir ineffective and whether excluding such genetic non-responders from the treatment group might help push it over the significance threshold.

Once imported into cells, remdesivir is gradually degraded to AMP analog, which is then converted to ATP analog to interfere with viral RdRp^33–35^. The same ATP analog may also compete with endogenous ATP to inhibit other cellular proteins, which may inflict damage. Importantly, such damage is enough to cause cell death in Eμ-Myc;Arf−/− cells (Fig. 1b), which allowed for a screen to discover genes crucial for remdesivir importation and NTP conversion. The two top hits in our remdesivir screen fell into such categories (Fig. 3b). We identified SLC29A3, a membrane transporter that facilitates the cellular uptake of chemical compounds. Among the more than 350 SLC proteins in the genome, only SLC29A3 was significantly enriched in our screen, suggesting its uniqueness for the importation of remdesivir. Our screen also identified HINT1 (Histidine Triad Nucleotide Binding Protein 1), which is an enzyme that breaks down phosphoramidate bonds on purine nucleotides. Remdesivir was designed such that once inside cells, it is converted to AMP analog in several steps^33^ (Fig. 3c). Importantly, one of which involves breakage of such a phosphoramidate bond. Our screen results suggested this step, catalyzed by a cellular enzyme HINT1, could be rate-limiting in generating active remdesivir metabolites inside cells. Taken together, our study indicated SLC29A3 and HINT1 could be two cellular proteins crucial for the cellular concentration of remdesivir and its conversion into active metabolites, pointing to the possibility that genetic and expression differences in these two genes may affect COVID-19 patients’ response to remdesivir treatment.

### Favipiravir

Favipiravir is an antiviral prodrug that partially resembles a purine base. Favipiravir exhibited potent antiviral activity in SARS-CoV-2 animal models^36^ and multiple clinical trials showed that favipiravir significantly shortened viral clearance time and sped up clinical recovery^11,37–39^. In our screen, the enriched sgRNAs in cells that survived favipiravir treatment were mostly targeting cell death genes (Fig. 3d). It is possible that multiple proteins may play redundant roles in the importation and/or activation of favipiravir, therefore the enriched sgRNA sets could not identify specific drug importers or metabolizing enzymes for favipiravir. However, the sgRNAs depleted from cells that survived favipiravir treatment indicated drug inactivation mechanisms, which are discussed below.

### Drug export and inactivation mechanisms

In cells that survived drug selection, we were also able to analyze which sgRNAs were strongly depleted after drug selection. Such sgRNAs point to genes whose deficiency enhances drug efficacy and/or toxicity. We first examined the ABC family of drug exporters ^40^. Such proteins pump chemicals out of cells, therefore their deficiency will increase cellular drug concentration and result in higher levels of death. We examined in our COVID-19 drug screens, sgRNAs against which ABC genes were the most prominently depleted (Fig. 4a and a’, Extended Data Table 1). The results suggested that ABCC1 is a candidate exporter for remdesivir, and ABCC1, ABCB1 for colchicine.

**Figure 4.**
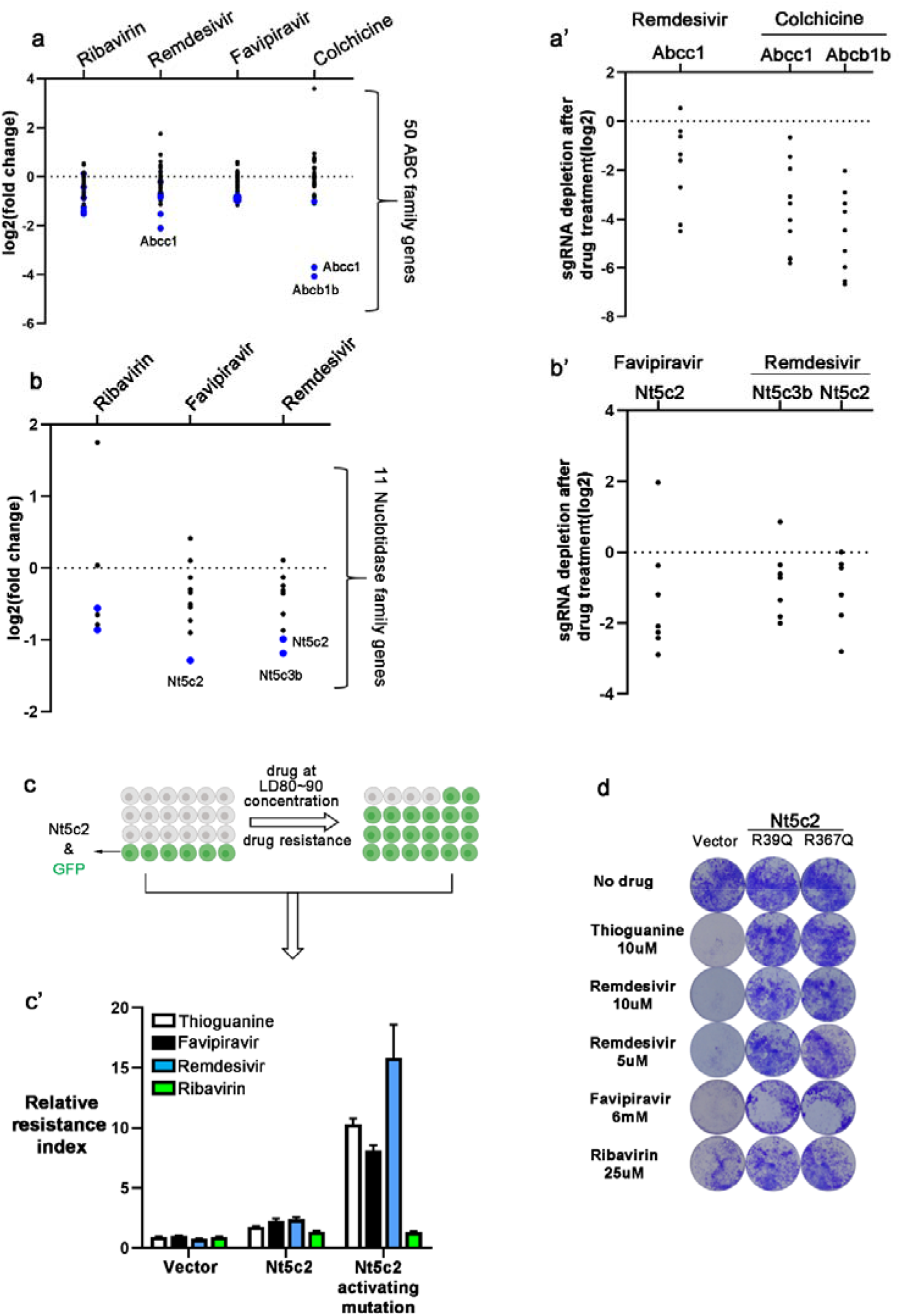
Analysis of the depleted gene sets with regards to ABC transporters and nucleotidases. **a)** and **b)** How genes belonging to the families of ABC transporters and nucleotidases behaved in the screens. Each dot represents a gene. Blue dots indicate genes whose depletion is statistically significant (p<0.05). **a’)** and **b’)** How individual sgRNA targeting the listed genes behaved in the drug screen. **c)** An experiment setting to assess whether NT5C2 activity protects cells from drug-induced toxicity. Eμ-Myc;Arf−/− cells partially infected by NT5C2 and GFP-expressing virus were subjected to drug treatment at concentrations that killed 80~90% (LD80-90) of parental cells. Drug inactivation by NT5C2 enabled better survival of NT5C2&GFP expressing cells, leading to increase of GFP positive rate in surviving cells. **c’)** Relative resistance index calculated from GFP percentages before and after drug treatment. Shown are Mean±SD from a triplicate experiment. Enhanced NT5C2 activity potently protected cells from remdesivir and favipiravir. Thioguanine was used as a positive control. **d)** Enhanced NT5C2 activity protected 293T cells from thioguanine, remdesivir and favipiravir.

As discussed before, the efficacy of antiviral drugs strongly depends on the cellular concentration of their respective NTP analogs. For nucleoside analogs, the first phosphorylation step to generate NMP analog is the rate-limiting step for activation. Consequently, nucleotidases that break down NMP analog could significantly reduce the antiviral efficacy of such drugs. Indeed, recent animal studies showed that for remdesivir, significant amounts of the AMP analog were dephosphorylated into inactive adenosine analog (depicted in Fig. 3c) ^34,41^, which could greatly reduce the antiviral efficacy of remdesivir. Since this type of dephosphorylation is carried out by 5’-Nucleotidases^42^, we examined all 5’-Nucleotidase genes in our screen (Extended Data Table 2) and found that NT5C3B and NT5C2 are potentially responsible for such a dephosphorylation reaction for remdesivir. A similar analysis also suggested that NT5C2 is responsible for the dephosphorylation of NMP analog for favipiravir (Fig. 4b and b’).

Notably, in the treatment of leukemia, NT5C2 activity is the most frequent cause of thiopurine treatment failure and disease relapse ^43^. Through dephosphorylation of GMP analog formed by thiopurine, this enzyme inactivates thiopurine. Based on our screen results, we asked whether NT5C2 could similarly inactivate remdesivir and favipiravir. Indeed, increased cellular NT5C2 activity significantly reduced cytotoxicity of both remdesivir and favipiravir in multiple cell lines (Fig. 4c and 4d), but had little effects on ribavirin. This suggested that analogous to the case of thiopurine, NT5C2 critically reduces the NMP analog of remdesivir and favipiravir, consequently reducing the level of active antiviral NTP analogs. Therefore, NT5C2 may also represent a crucial point for determining the antiviral efficacy of remdesivir and favipiravir.

### Validation of sgRNA screen results

Our screens, leveraging NTP analog-induced cell death, discovered genes that affect the import, export and metabolism of antiviral drugs used in COVID-19 treatment. Prompted by the observation with NT5C2, we further validated top hits for three antiviral drugs in our study (Fig. 5a). Consistent with the screen data, sgRNAs targeting SLC29A3 and HINT1 significantly protected cells from remdesivir, whereas sgRNAs targeting ABCC1, NT5C2 and NT5C3B caused sensitization. Similar validation results were also seen for genes discovered in favipiravir and ribavirin screens, highlighting their important roles in regulating the efficacy of these drugs.

**Figure 5.**
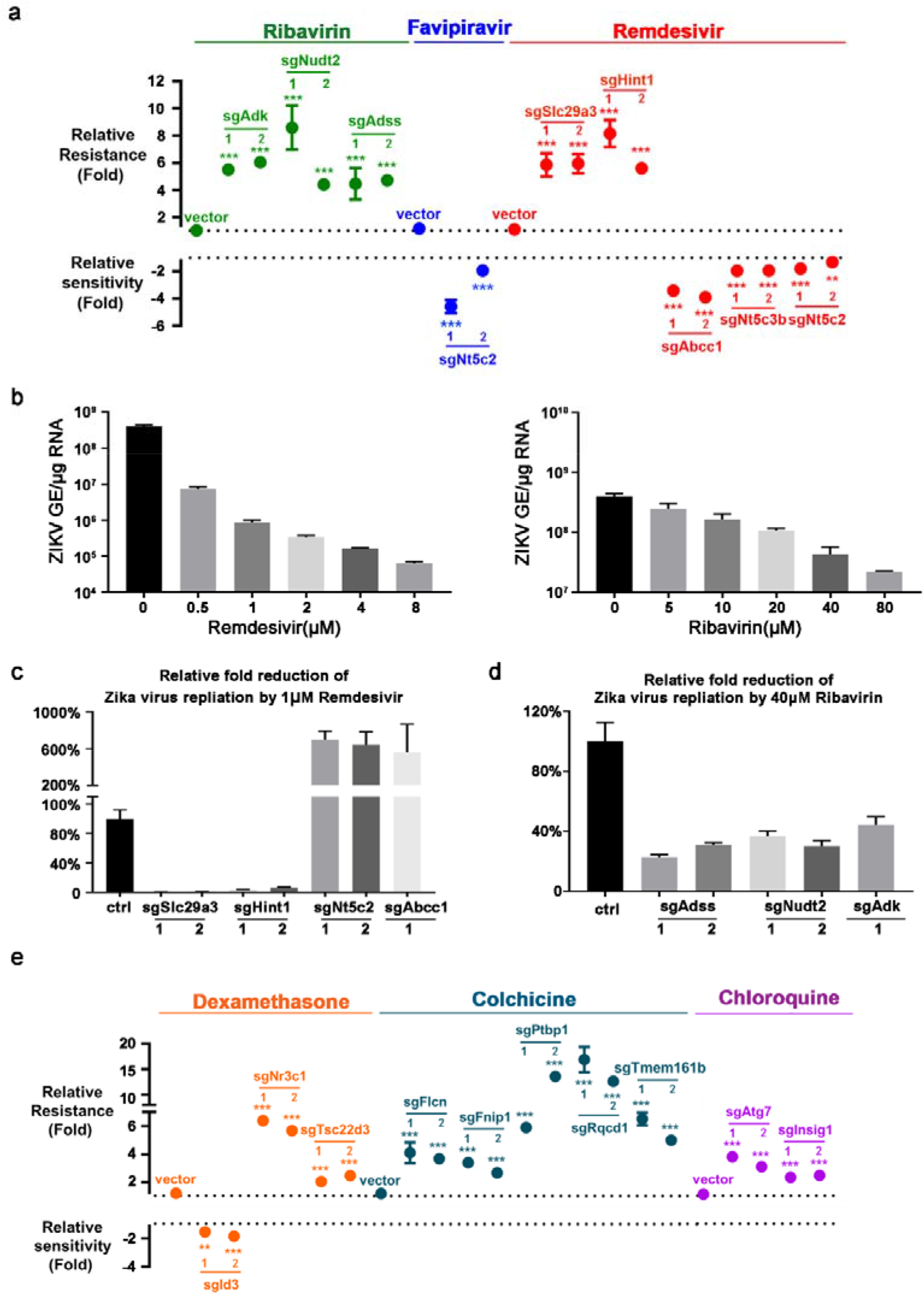
Validation of CRISPR screen results for COVID-19 drugs. **a and e)**. Effects of indicated screen hit genes on drug sensitivity. sgRNAs targeting the indicated genes were expressed in Eμ-Myc;Arf−/− cells. An experimental setting similar to Fig.4c was used to assess whether a gene knockout caused resistance or sensitization to COVID-19 drugs. Shown are Mean±SEM from three independent experiments. **p<0.05. ***p<0.01. **b)** Remdesivir and ribavirin suppressed Zika virus replication in Huh7 cells. Antiviral efficacies of remdesivir **(c)** and ribavirin **(d)** were impacted by genes identified in CRISPR screens. Shown are Mean±SD from a triplet experiment.

Due to lack of timely access, we could not further confirm our finding with Sars-CoV-2 models. As an alternative, we used Zika virus, whose replication^44^ could be suppressed by remdesivir and ribavirin (Fig. 5b). We knocked out SLC29A3, HINT1, ABCC1 and NT5C2 in Huh7 cells and found that deficiencies of these genes indeed significantly affected remdesivir’s antiviral efficacy (Fig. 5c). For example, HINT1 or SLC29A3 deficiency reduced remdesivir’s antiviral efficacy by 30 to 100 folds. This demonstrated the essential roles of SLC29A3 and HINT1 in the importation and activation of remdesivir. Conversely, knockout of ABCC1 or NT5C2 caused significant enhancement of remdesivir’s antiviral efficacy, confirming their importance as the exporter and inactivator of remdesivir.

In the case of ribavirin, knockout of genes identified in this study, including ADK; ADSS and NUDT2 also decreased ribavirin’s antiviral efficacy (Fig. 5d). Taken together, these results confirmed the virological relevance of our screen strategy and provided the first set of potential genetic biomarkers for remdesivir.

Lastly, we also cloned sgRNAs to interrogate some of the top hits in colchicine and dexamethasone screens. We confirmed that knockout of such genes indeed conferred resistance or hypersensitivity to corresponding drugs (Fig. 5e), indicating they may modulate the immunosuppressive effects of these drugs. For example, TMEM161D, RQCD1, PTBP1, FLCN and FNIP knockout significantly protected cells from colchicine-induced death. Our results for the first time linked these genes to microtubule poisons. We also confirmed that ID3 knockout sensitized cells to dexamethasone. Taken together, screens conducted in this study will help advance the understanding of the action of COVID-19 drugs, potentially pointing to genetic biomarkers for disease management.

## Discussion

Dexamethasone, colchicine, remdesivir, favipiravir and ribavirin have been shown to speed up SARS-coV-2 viral clearance or reduce COVID-19 death rate by various studies. The role of chloroquine and hydroxychloroquine in COVID-19 is more controversial^13–16,45–47^, with larger studies of hospitalized patients arguing against their use^14–16^. We also conducted CRISPR screens on chloroquine phosphate and hydroxychloroquine and identified nearly identical lists of resistance genes (Extended Data Fig. 2). Our results suggest that a group of genes involved in autophagy, as well as lipid and cholesterol synthesis significantly modulate the action of chloroquine drugs. Such a gene list may be useful for understanding chloroquine drugs’ clinical use.

Due to the urgency of COVID-19 situation and lack of relevant knowledge, COVID-19 drugs have been applied with little consideration of genetic biomarkers. Given the rapid disease progression of COVID-19 and the presence of multiple treatment options, it is urgent to discover genetic modulators for each COVID-19 drug, such that patients may be matched to more effective drug choice. Here we present the first report on genome-wide CRISPR screens on these COVID-19 drugs, with data on both resistance and sensitization mechanisms.

For antiviral drugs, low toxicity and lack of executable screen endpoints presented two significant hurdles for genome-wide screens. Our approach, using NTP analog-induced cell death as a surrogate screen endpoint, overcame these obstacles and helped uncover the transport and metabolism mechanisms crucial for the antiviral efficacy of these drugs. From a method standpoint, this screen strategy is the first that is capable of employing genome-wide libraries to identify genetic biomarkers for antiviral drugs. Such a screen platform is broadly applicable for other types of nucleoside analogs used to treat existed and emerging viral diseases. Extended Data Figure 3 summarized our findings and showed the relative uniqueness of the drug transporters and metabolizing enzymes discovered for these COVID-19 drugs.

The clinical efficacies of several COVID-19 drugs including remdesivir remain under debate^48^. Incorporation of genetic biomarkers of drugs may lead to better assessment of their efficacy, as demonstrated by previous clinical trials of PARP inhibitors^49^. Importantly, for nucleoside-based anticancer drugs that are structurally similar to antiviral drugs, numerous clinical studies have shown that SNPs and expression status of their relevant transporters and metabolizing enzymes dictated clinical outcome^50–54^. Therefore, the transportation, activating and detoxifying mechanisms uncovered in this screen may point to clinical biomarkers of COVID-19 drugs. If such knowledge could be established in the future, it may help better understand the efficacy of individual drugs in the context of COVID-19. For example, a SNP on NT5C2, present in 15% of the population, significantly affected the cellular activity of thiopurine and its incorporation into DNA^55^. Our results would similarly suggest that patients with such genetic variation may respond differently to remdesivir and favipiravir, but not to ribavirin (Fig. 4c and 4d). In addition, medically relevant SNPs for many of the genes identified in our screen have been reported in human population. For example, deleterious SNPs of SLC29A3, which we discovered as the transporter for remdesivir, underly a diverse range of disease phenotypes including short stature, osteoporosis, diabetes, hyperpigmentation, chronic inflammation and hearing loss^56–59^. Given our finding that SLC29A3 deficiency greatly reduced remdesivir’s antiviral effects (Fig. 5c), such a genetic background may call for further consideration on the use of remdesivir, yet the efficacy of favipiravir and ribavirin may be less affected. From a precision medicine perspective, our findings may provide rationales for pairing COVID-19 patients with proper drug choice, potentially improving the overall cure rate. Such knowledge may also be useful for the management of other diseases that utilize these drugs, such as influenza, Dengue and Zika virus infections.

Lastly, rational design of drug combination is still lacking for COVID-19 treatment. Our study may also shed light on such drug interaction. For example, given that NT5C2 causes the inactivation of both remdesivir and favipiravir, combinatorial use of both drugs may potentially saturate NT5C2 activity, which might lead to better antiviral outcome for these drugs.

There are several limitations to our study. As a loss of function screen, our study cannot identify certain kinds of genes. For example, if several proteins function redundantly in a drug-metabolizing step, our screen will not be able to uncover such proteins either in the enriched or depleted sgRNA dataset. Genome-wide overexpression screens may help complement this study. Moreover, there are many discussions that drug efficacy in COVID-19 patients may be significantly influenced by the timing of treatment. Our screen cannot address that question either. Due to the urgency of the situation, we also did not have time to further delve into the mechanisms of some of the screen hits. We present our screen results in the current form to facilitate relevant research, in the hope that among the list of screen hits, drug biomarkers will be established, such that better understanding of COVID-19 treatment will be achieved to help combat the pandemic.

## Acknowledgement

We thank Drs. Dangsheng Li, Ting Ni, Yinan Zhang and Xin Hu for discussions and advice on this project and technical help from the Core Facility for Cell Biology and Core Facility for Molecular Biology at SIBCB. We thank MedChemExpress for kindly providing free samples of remdesivir for this study. This work was supported by the National Science and Technology Major Project of the Ministry of Science and Technology of China (Grant No. 2017YFA0504503; Grant No. 2018ZX10101004003001), the Strategic Priority Research Program of Chinese Academy of Sciences, Grant No. XDB19000000, Collaborative project of National Natural Science Foundation of China and Deutsche Forschungsgemeinschaft (81761138046), Natural Science Foundation of the People’s Republic of China (Grant No. 81972600), CAS Interdisciplinary Innovation Team Fund (JCTD-2018-14) and a Director’s Special Research Fund from SIBCB.

## Author Contributions

H.J. conceived, supervised the study and wrote the manuscript. W.J. performed the screens and validation. A.Y. established the screen platform and participated in validation experiments in this study. J.M. analyzed screen data; prepared figures and participated in the earlier steps of the screen platform, including transferring the genome-wide sgRNA library into a retroviral backbone. A.Y., J.M. and W.J. completed anticancer drugs screens referenced in this study. The Zika virus validation experiments were performed by D. L. under the supervision of G. L. L. X., M. L., C. W., Z. H., S. C., J. Z., S.C., Q. J., Y.C. and L.S. participated in drug screens or provided technical helps to establish the screen protocol. Z. Z. and Y. Z. advised on and provided helps to the screen platform.

## Data and material availability

All data is available in the main text or the supplementary materials. Plasmids generated in this study will be deposited to the plasmid sharing platform at BIO-Research Innovation Center Suzhou (www.brics.ac.cn/plasmid/?columnId=15) for distribution.

---

**Extended Data Figure 1.**
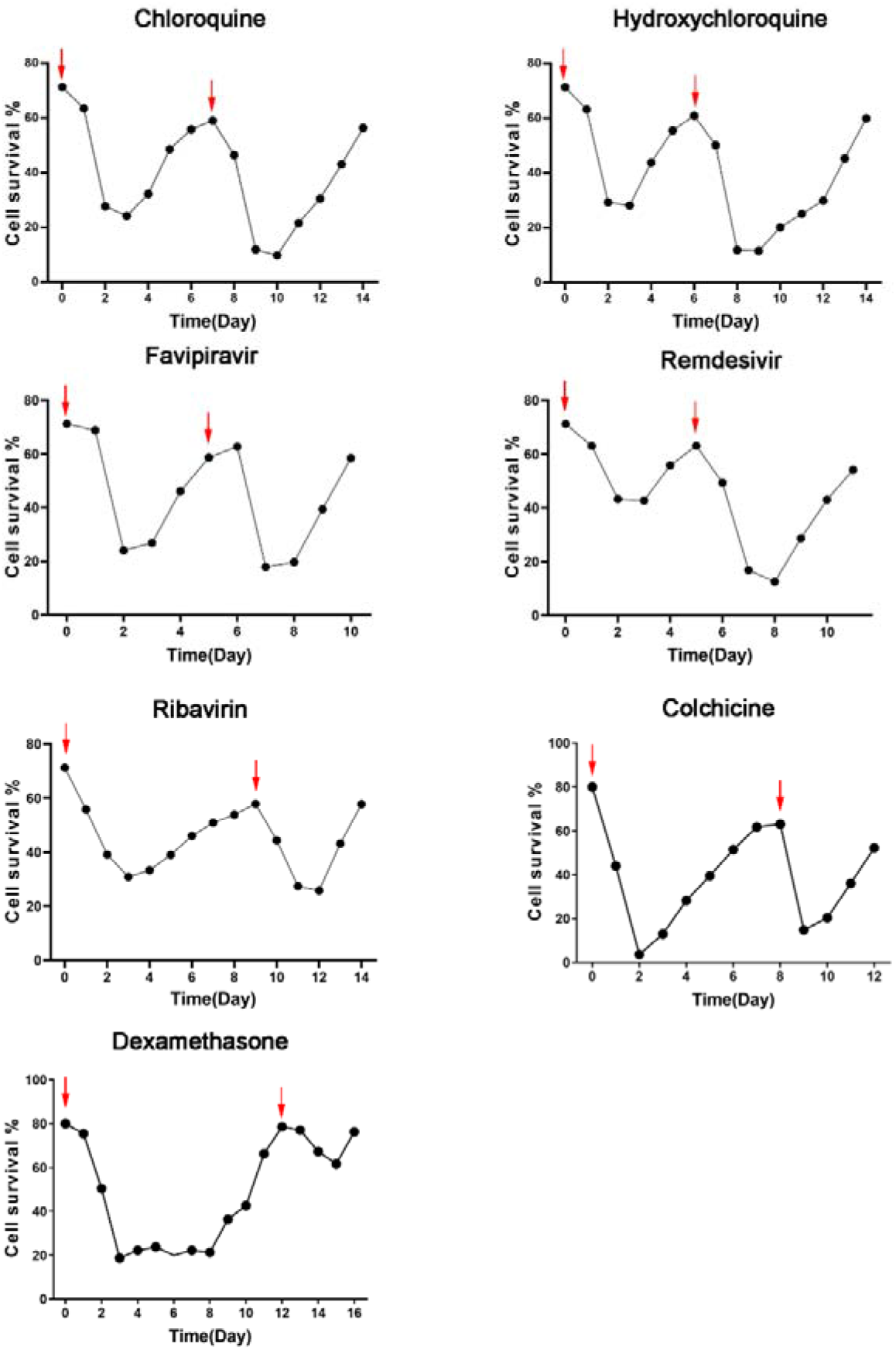
Survival curves of cells during COVID-19 drug screens. Eμ-Myc;Arf−/− cells were subjected to two rounds of treatment with the indicated drugs. Overall cellular viabilities were analyzed every day during the duration of the experiment. Blue arrows indicated the date when drugs were applied. Drugs were used at the following concentrations: remdesivir (16.4μM), ribavirin (20μM), favipiravir (400μM), chloroquine diphosphate (10μM), hydroxychloroquine sulfate (11.2μM), colchicine (25nM) and dexamethasone (4.7μM).

**Extended Data Figure 2.**
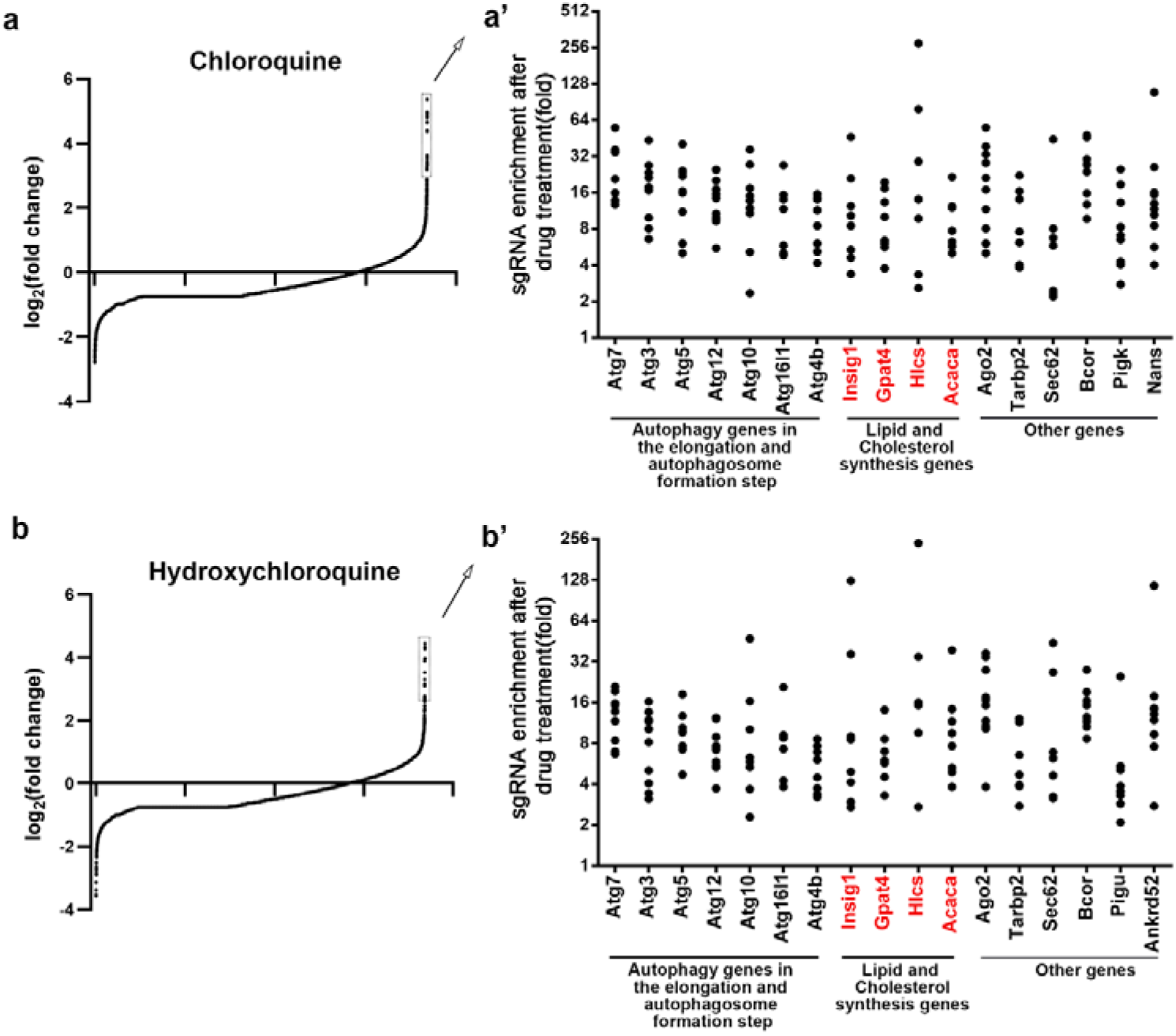
Genes enriched in chloroquine and hydroxychloroquine screens. **a and b)** Figures showing the overall distribution patterns of genes in the drug screen. Top enriched genes are boxed out and detailed in **a’ and b’)**, which show the extent of enrichment of individual sgRNAs after drug selection. In **a’ and b’)**, each dot represents a sgRNA targeting the indicated gene. p values for all genes depicted in **a’ and b’)** are lower than 0.0002.

**Extended Data Figure 3.**
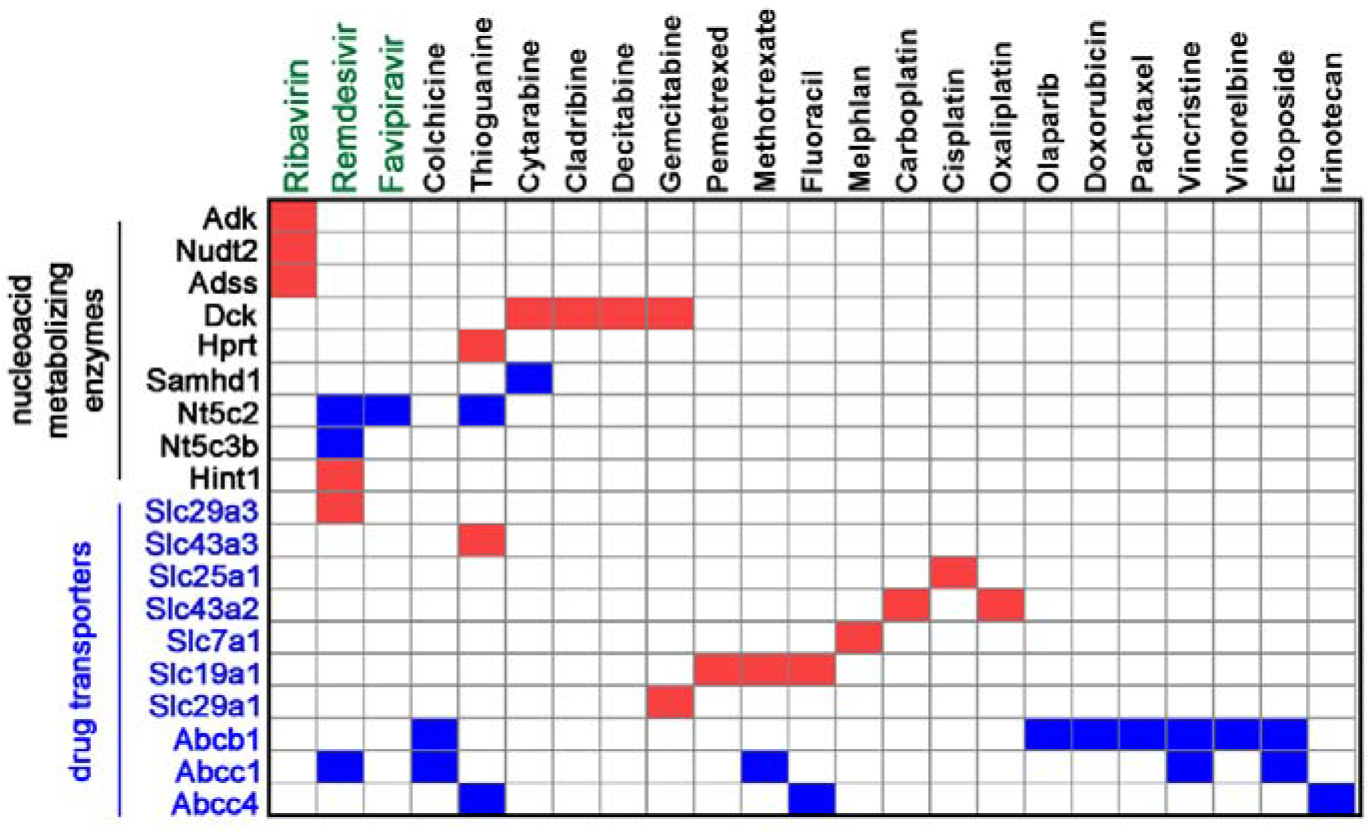
Summary and comparison of drug metabolizing enzymes and transporters identified in genome-wide CRISPR screens using our platform. Red boxes indicate enrichment of sgRNA against a gene in corresponding drug screen, whereas blue boxes indicate depletion.

**Extended Dataset Table 1.**
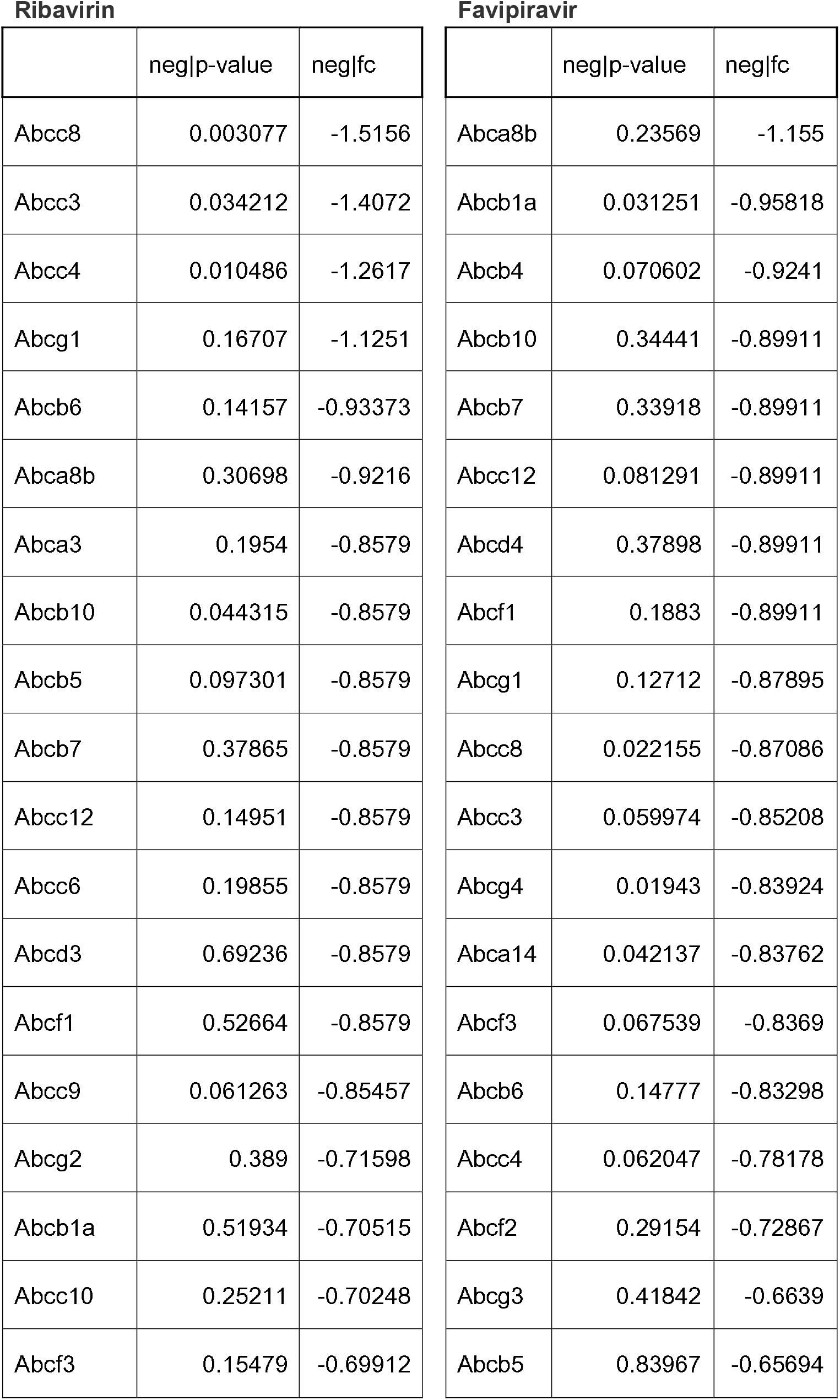
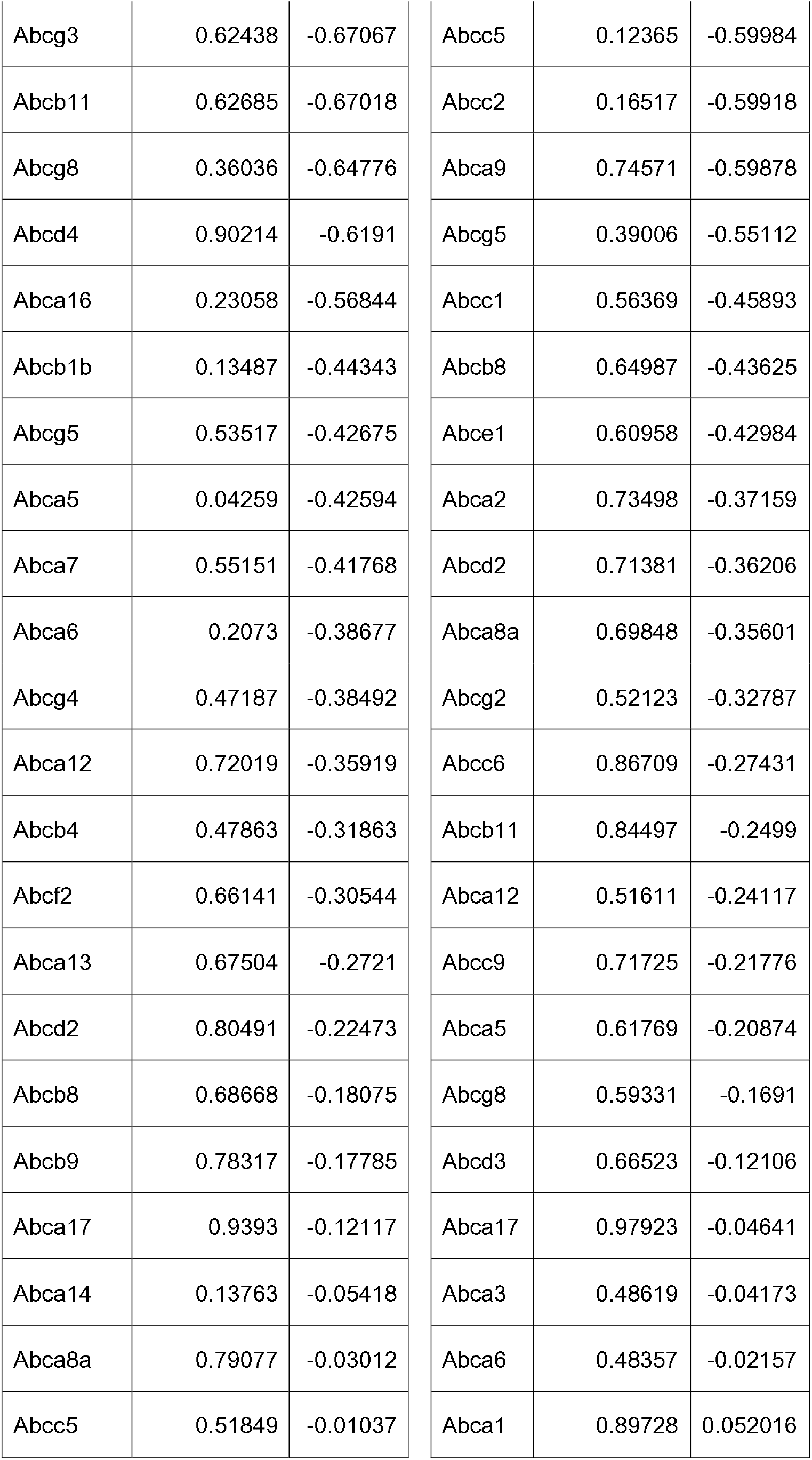
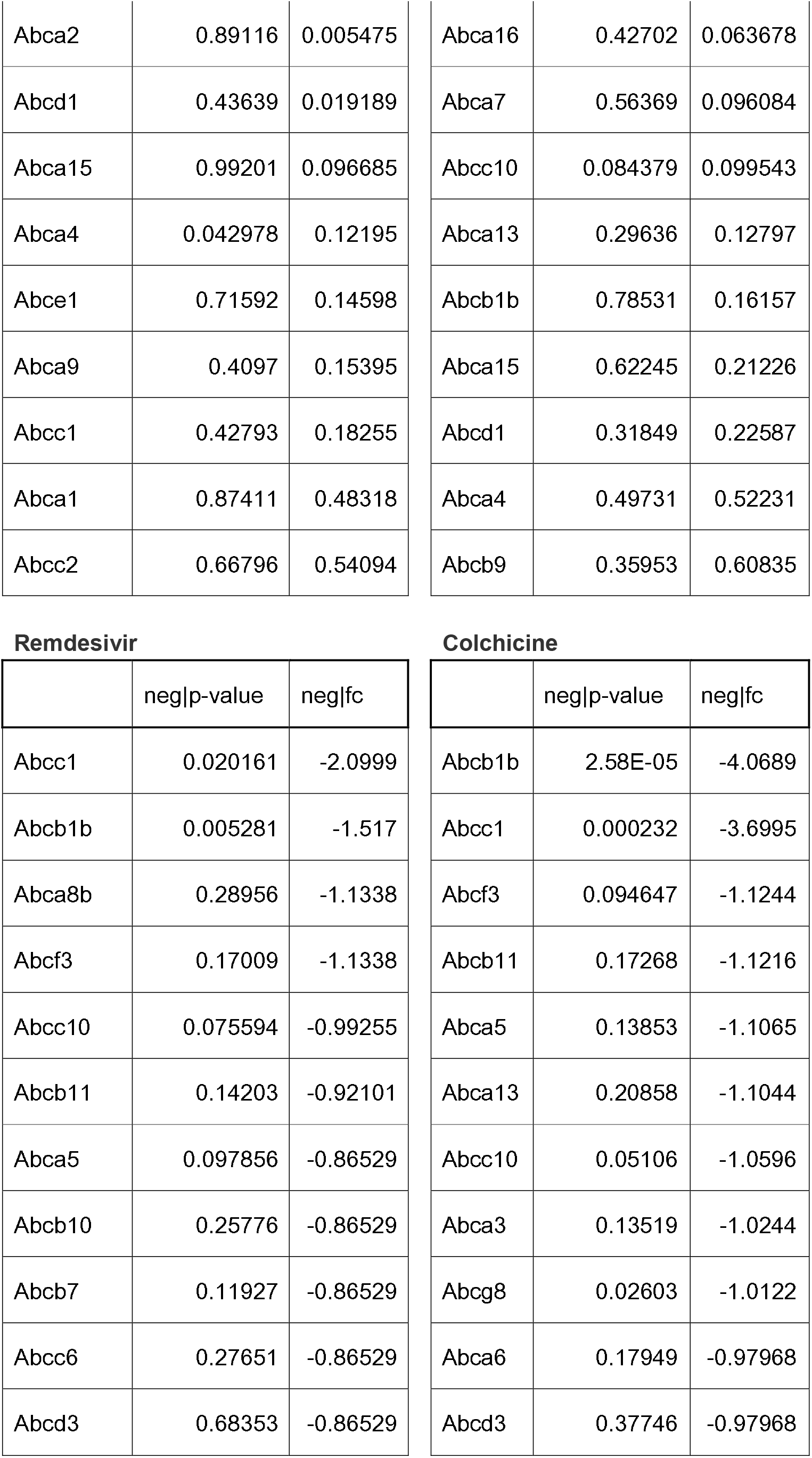
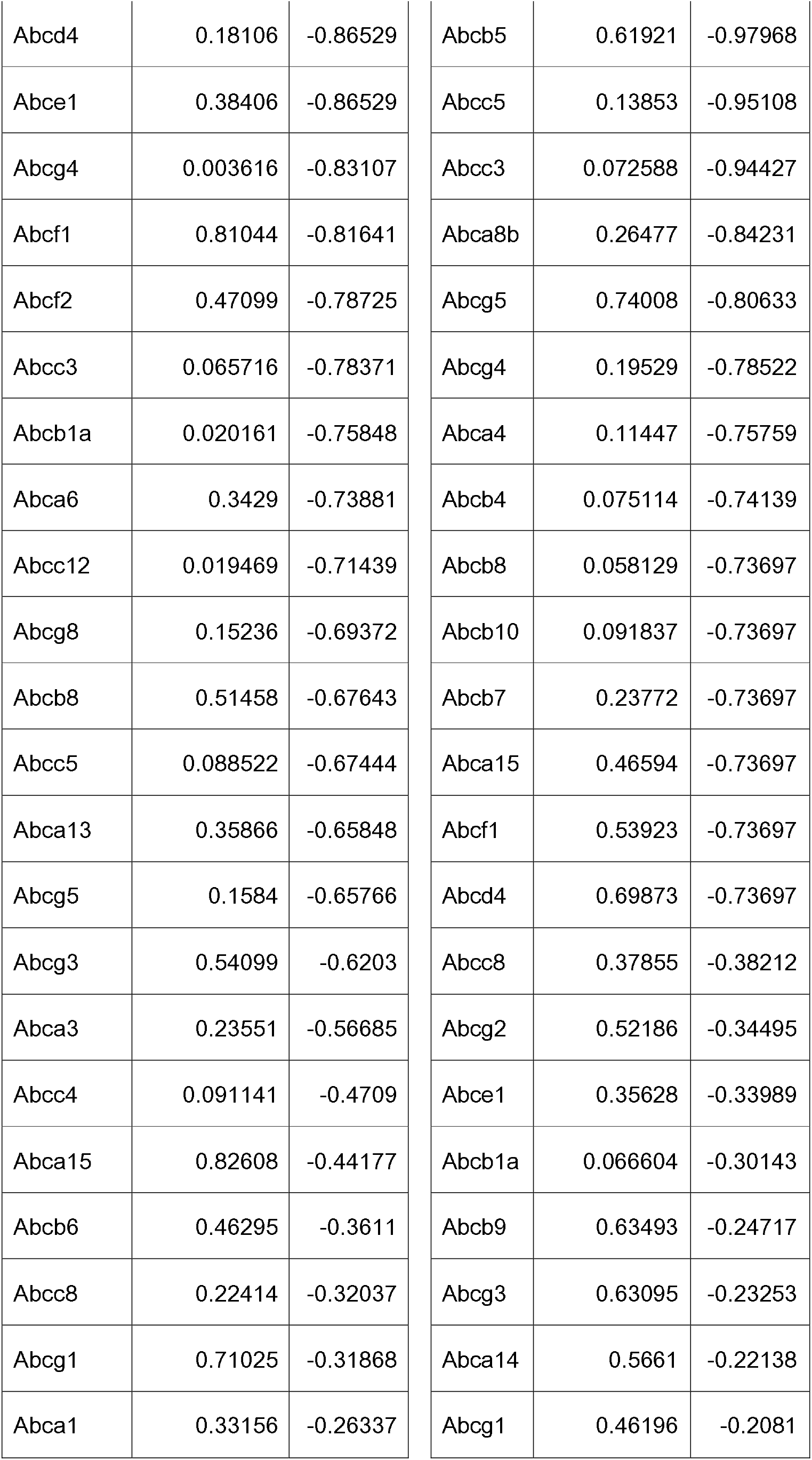
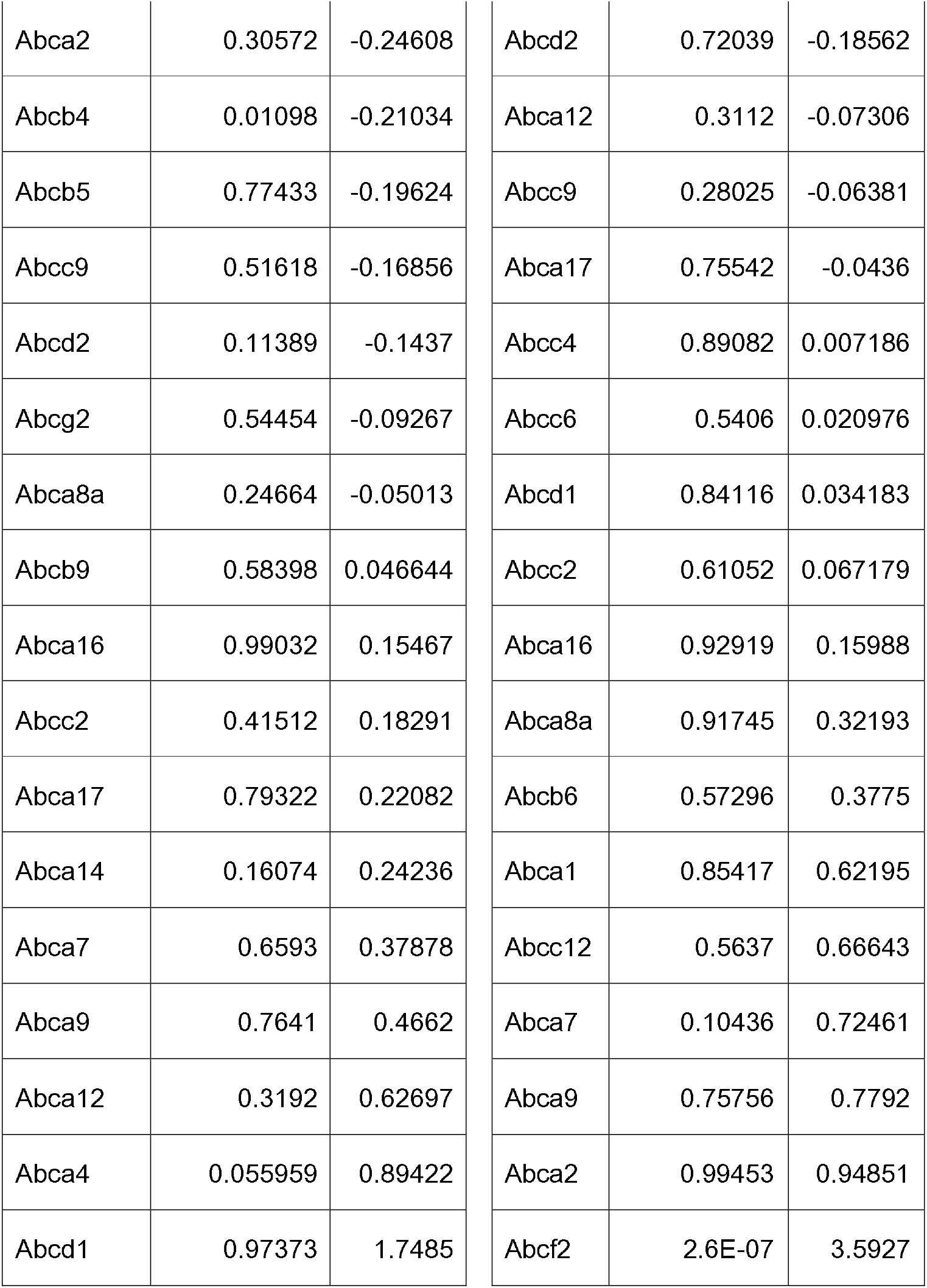
sgRNA depletion data of ABC family genes.

| **Ribavirin** |  |  | **Favipiravir** |  |  |
| --- | --- | --- | --- | --- | --- |
|  | neg\|p-value | neg\|fc |  | neg\|p-value | neg\|fc |
| Abcc8 | 0.003077 | -1.5156 | Abca8b | 0.23569 | -1.155 |
| Abcc3 | 0.034212 | -1.4072 | Abcb1a | 0.031251 | -0.95818 |
| Abcc4 | 0.010486 | -1.2617 | Abcb4 | 0.070602 | -0.9241 |
| Abcg1 | 0.16707 | -1.1251 | Abcb10 | 0.34441 | -0.89911 |
| Abcb6 | 0.14157 | -0.93373 | Abcb7 | 0.33918 | -0.89911 |
| Abca8b | 0.30698 | -0.9216 | Abcc12 | 0.081291 | -0.89911 |
| Abca3 | 0.1954 | -0.8579 | Abcd4 | 0.37898 | -0.89911 |
| Abcb10 | 0.044315 | -0.8579 | Abcf1 | 0.1883 | -0.89911 |
| Abcb5 | 0.097301 | -0.8579 | Abcg1 | 0.12712 | -0.87895 |
| Abcb7 | 0.37865 | -0.8579 | Abcc8 | 0.022155 | -0.87086 |
| Abcc12 | 0.14951 | -0.8579 | Abcc3 | 0.059974 | -0.85208 |
| Abcc6 | 0.19855 | -0.8579 | Abcg4 | 0.01943 | -0.83924 |
| Abcd3 | 0.69236 | -0.8579 | Abca14 | 0.042137 | -0.83762 |
| Abcf1 | 0.52664 | -0.8579 | Abcf3 | 0.067539 | -0.8369 |
| Abcc9 | 0.061263 | -0.85457 | Abcb6 | 0.14777 | -0.83298 |
| Abcg2 | 0.389 | -0.71598 | Abcc4 | 0.062047 | -0.78178 |
| Abcb1a | 0.51934 | -0.70515 | Abcf2 | 0.29154 | -0.72867 |
| Abcc10 | 0.25211 | -0.70248 | Abcg3 | 0.41842 | -0.6639 |
| Abcf3 | 0.15479 | -0.69912 | Abcb5 | 0.83967 | -0.65694 |
| Abcg3 | 0.62438 | -0.67067 | Abcc5 | 0.12365 | -0.59984 |
| Abcb11 | 0.62685 | -0.67018 | Abcc2 | 0.16517 | -0.59918 |
| Abcg8 | 0.36036 | -0.64776 | Abca9 | 0.74571 | -0.59878 |
| Abcd4 | 0.90214 | -0.6191 | Abcg5 | 0.39006 | -0.55112 |
| Abca16 | 0.23058 | -0.56844 | Abcc1 | 0.56369 | -0.45893 |
| Abcb1b | 0.13487 | -0.44343 | Abcb8 | 0.64987 | -0.43625 |
| Abcg5 | 0.53517 | -0.42675 | Abce1 | 0.60958 | -0.42984 |
| Abca5 | 0.04259 | -0.42594 | Abca2 | 0.73498 | -0.37159 |
| Abca7 | 0.55151 | -0.41768 | Abcd2 | 0.71381 | -0.36206 |
| Abca6 | 0.2073 | -0.38677 | Abca8a | 0.69848 | -0.35601 |
| Abcg4 | 0.47187 | -0.38492 | Abcg2 | 0.52123 | -0.32787 |
| Abca12 | 0.72019 | -0.35919 | Abcc6 | 0.86709 | -0.27431 |
| Abcb4 | 0.47863 | -0.31863 | Abcb11 | 0.84497 | -0.2499 |
| Abcf2 | 0.66141 | -0.30544 | Abca12 | 0.51611 | -0.24117 |
| Abca13 | 0.67504 | -0.2721 | Abcc9 | 0.71725 | -0.21776 |
| Abcd2 | 0.80491 | -0.22473 | Abca5 | 0.61769 | -0.20874 |
| Abcb8 | 0.68668 | -0.18075 | Abcg8 | 0.59331 | -0.1691 |
| Abcb9 | 0.78317 | -0.17785 | Abcd3 | 0.66523 | -0.12106 |
| Abca17 | 0.9393 | -0.12117 | Abca17 | 0.97923 | -0.04641 |
| Abca14 | 0.13763 | -0.05418 | Abca3 | 0.48619 | -0.04173 |
| Abca8a | 0.79077 | -0.03012 | Abca6 | 0.48357 | -0.02157 |
| Abcc5 | 0.51849 | -0.01037 | Abca1 | 0.89728 | 0.052016 |
| Abca2 | 0.89116 | 0.005475 | Abca16 | 0.42702 | 0.063678 |
| Abcd1 | 0.43639 | 0.019189 | Abca7 | 0.56369 | 0.096084 |
| Abca15 | 0.99201 | 0.096685 | Abcc10 | 0.084379 | 0.099543 |
| Abca4 | 0.042978 | 0.12195 | Abca13 | 0.29636 | 0.12797 |
| Abce1 | 0.71592 | 0.14598 | Abcb1b | 0.78531 | 0.16157 |
| Abca9 | 0.4097 | 0.15395 | Abca15 | 0.62245 | 0.21226 |
| Abcc1 | 0.42793 | 0.18255 | Abcd1 | 0.31849 | 0.22587 |
| Abca1 | 0.87411 | 0.48318 | Abca4 | 0.49731 | 0.52231 |
| Abcc2 | 0.66796 | 0.54094 | Abcb9 | 0.35953 | 0.60835 |

Remdesivir
|  | neg\|p-value | neg\|fc |
| --- | --- | --- |
| Abcc1 | 0.020161 | -2.0999 |
| Abcb1b | 0.005281 | -1.517 |
| Abca8b | 0.28956 | -1.1338 |
| Abcf3 | 0.17009 | -1.1338 |
| Abcc10 | 0.075594 | -0.99255 |
| Abcb11 | 0.14203 | -0.92101 |
| Abca5 | 0.097856 | -0.86529 |
| Abcb10 | 0.25776 | -0.86529 |
| Abcb7 | 0.11927 | -0.86529 |
| Abcc6 | 0.27651 | -0.86529 |
| Abcd3 | 0.68353 | -0.86529 |

Colchicine
|  | neg\|p-value | neg\|fc |
| --- | --- | --- |
| Abcb1b | 2.58E-05 | -4.0689 |
| Abcc1 | 0.000232 | -3.6995 |
| Abcf3 | 0.094647 | -1.1244 |
| Abcb11 | 0.17268 | -1.1216 |
| Abca5 | 0.13853 | -1.1065 |
| Abca13 | 0.20858 | -1.1044 |
| Abcc10 | 0.05106 | -1.0596 |
| Abca3 | 0.13519 | -1.0244 |
| Abcg8 | 0.02603 | -1.0122 |
| Abca6 | 0.17949 | -0.97968 |
| Abcd3 | 0.37746 | -0.97968 |
| Abcd4 | 0.18106 | -0.86529 | Abcb5 | 0.61921 | -0.97968 |
| Abce1 | 0.38406 | -0.86529 | Abcc5 | 0.13853 | -0.95108 |
| Abcg4 | 0.003616 | -0.83107 | Abcc3 | 0.072588 | -0.94427 |
| Abcf1 | 0.81044 | -0.81641 | Abca8b | 0.26477 | -0.84231 |
| Abcf2 | 0.47099 | -0.78725 | Abcg5 | 0.74008 | -0.80633 |
| Abcc3 | 0.065716 | -0.78371 | Abcg4 | 0.19529 | -0.78522 |
| Abcb1a | 0.020161 | -0.75848 | Abca4 | 0.11447 | -0.75759 |
| Abca6 | 0.3429 | -0.73881 | Abcb4 | 0.075114 | -0.74139 |
| Abcc12 | 0.019469 | -0.71439 | Abcb8 | 0.058129 | -0.73697 |
| Abcg8 | 0.15236 | -0.69372 | Abcb10 | 0.091837 | -0.73697 |
| Abcb8 | 0.51458 | -0.67643 | Abcb7 | 0.23772 | -0.73697 |
| Abcc5 | 0.088522 | -0.67444 | Abca15 | 0.46594 | -0.73697 |
| Abca13 | 0.35866 | -0.65848 | Abcf1 | 0.53923 | -0.73697 |
| Abcg5 | 0.1584 | -0.65766 | Abcd4 | 0.69873 | -0.73697 |
| Abcg3 | 0.54099 | -0.6203 | Abcc8 | 0.37855 | -0.38212 |
| Abca3 | 0.23551 | -0.56685 | Abcg2 | 0.52186 | -0.34495 |
| Abcc4 | 0.091141 | -0.4709 | Abce1 | 0.35628 | -0.33989 |
| Abca15 | 0.82608 | -0.44177 | Abcb1a | 0.066604 | -0.30143 |
| Abcb6 | 0.46295 | -0.3611 | Abcb9 | 0.63493 | -0.24717 |
| Abcc8 | 0.22414 | -0.32037 | Abcg3 | 0.63095 | -0.23253 |
| Abcg1 | 0.71025 | -0.31868 | Abca14 | 0.5661 | -0.22138 |
| Abca1 | 0.33156 | -0.26337 | Abcg1 | 0.46196 | -0.2081 |
| Abca2 | 0.30572 | -0.24608 | Abcd2 | 0.72039 | -0.18562 |
| Abcb4 | 0.01098 | -0.21034 | Abca12 | 0.3112 | -0.07306 |
| Abcb5 | 0.77433 | -0.19624 | Abcc9 | 0.28025 | -0.06381 |
| Abcc9 | 0.51618 | -0.16856 | Abca17 | 0.75542 | -0.0436 |
| Abcd2 | 0.11389 | -0.1437 | Abcc4 | 0.89082 | 0.007186 |
| Abcg2 | 0.54454 | -0.09267 | Abcc6 | 0.5406 | 0.020976 |
| Abca8a | 0.24664 | -0.05013 | Abcd1 | 0.84116 | 0.034183 |
| Abcb9 | 0.58398 | 0.046644 | Abcc2 | 0.61052 | 0.067179 |
| Abca16 | 0.99032 | 0.15467 | Abca16 | 0.92919 | 0.15988 |
| Abcc2 | 0.41512 | 0.18291 | Abca8a | 0.91745 | 0.32193 |
| Abca17 | 0.79322 | 0.22082 | Abcb6 | 0.57296 | 0.3775 |
| Abca14 | 0.16074 | 0.24236 | Abca1 | 0.85417 | 0.62195 |
| Abca7 | 0.6593 | 0.37878 | Abcc12 | 0.5637 | 0.66643 |
| Abca9 | 0.7641 | 0.4662 | Abca7 | 0.10436 | 0.72461 |
| Abca12 | 0.3192 | 0.62697 | Abca9 | 0.75756 | 0.7792 |
| Abca4 | 0.055959 | 0.89422 | Abca2 | 0.99453 | 0.94851 |
| Abcd1 | 0.97373 | 1.7485 | Abcf2 | 2.6E-07 | 3.5927 |

**Extended Dataset Table 2.** sgRNA depletion data of NT5 family genes.

**Ribavirin**
|  | neg\|p-value | neg\|fc |
| --- | --- | --- |
| Nt5c | 0.0048494 | -0.8579 |
| Nt5c1b | 0.59428 | -0.8579 |
| Nt5c3 | 0.46134 | -0.8579 |
| Nt5c3b | 0.18998 | -0.78704 |
| Nt5dc2 | 0.36552 | -0.65088 |
| Nt5c1a | 0.097767 | -0.56254 |
| Nt5dc1 | 0.46624 | -0.55633 |
| Nt5dc3 | 0.39761 | 0.038616 |
| Nt5m | 0.85146 | 0.03982 |
| Nt5e | 0.95324 | 0.043502 |
| Nt5c2 | 0.71592 | 1.7507 |

**Remdesivir**
|  | neg\|p-value | neg\|fc |
| --- | --- | --- |
| Nt5c3b | 0.0084289 | -1.1827 |
| Nt5c2 | 0.032509 | -0.98832 |
| Nt5c | 0.26641 | -0.86529 |
| Nt5c3 | 0.35866 | -0.86529 |
| Nt5c1b | 0.64403 | -0.63843 |
| Nt5dc1 | 0.32554 | -0.34892 |
| Nt5c1a | 0.80472 | -0.31396 |
| Nt5dc2 | 0.10876 | -0.31347 |
| Nt5dc3 | 0.54549 | -0.24357 |
| Nt5e | 0.77701 | -0.12608 |
| Nt5m | 0.95614 | 0.11014 |

**Favipiravir**
|  | neg\|p-value | neg\|fc |
| --- | --- | --- |
| Nt5c2 | 0.0002462 | -1.284 |
| Nt5c | 0.38607 | -0.89911 |
| Nt5dc2 | 0.26132 | -0.72846 |
| Nt5c1b | 0.58077 | -0.54048 |
| Nt5c1a | 0.14884 | -0.50111 |
| Nt5c3b | 0.34501 | -0.3387 |
| Nt5c3 | 0.2497 | -0.30023 |
| Nt5m | 0.84726 | -0.13039 |
| Nt5dc1 | 0.88353 | 0.10464 |
| Nt5e | 0.6698 | 0.41373 |
| Nt5dc3 | 0.59289 | 0.41744 |

**Extended Dataset Table 3.**
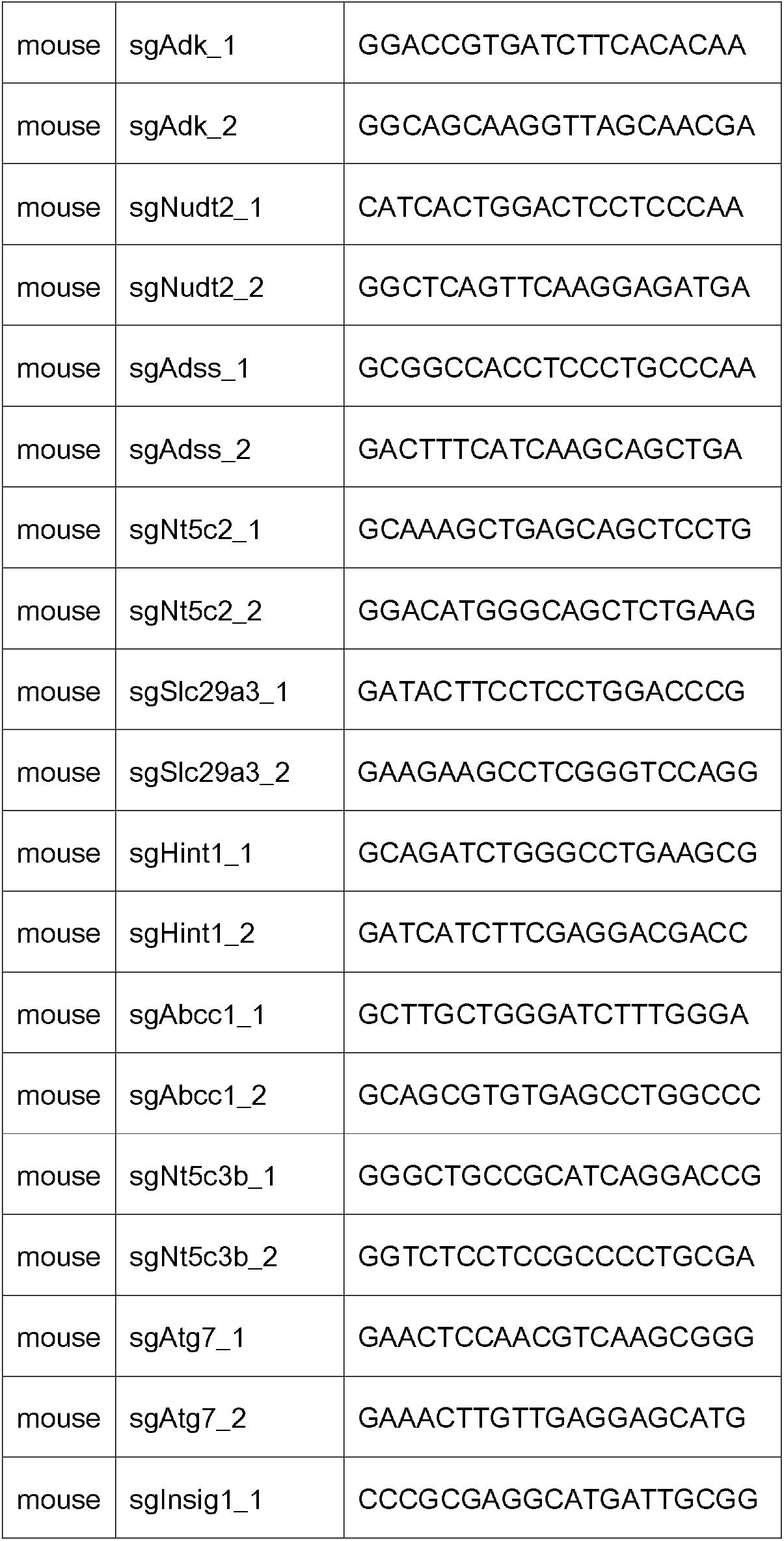
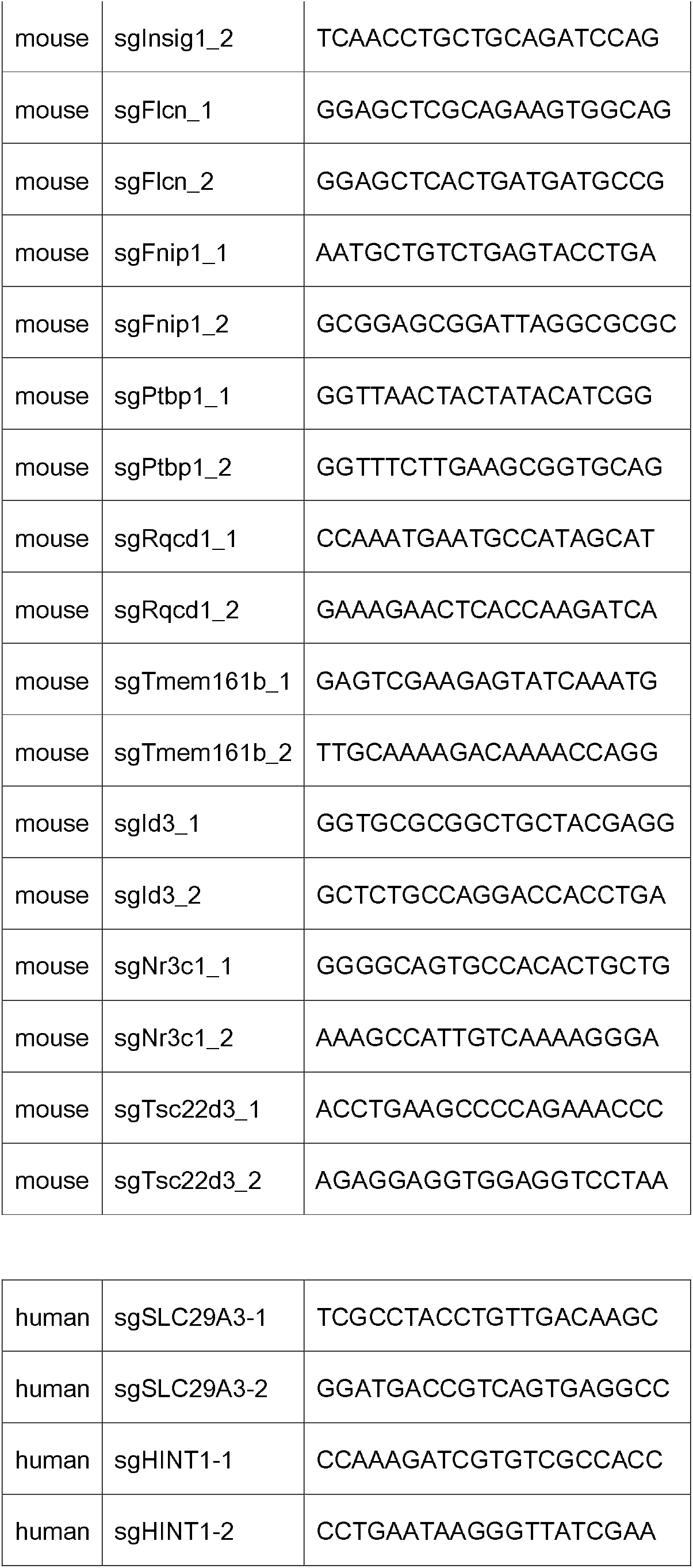
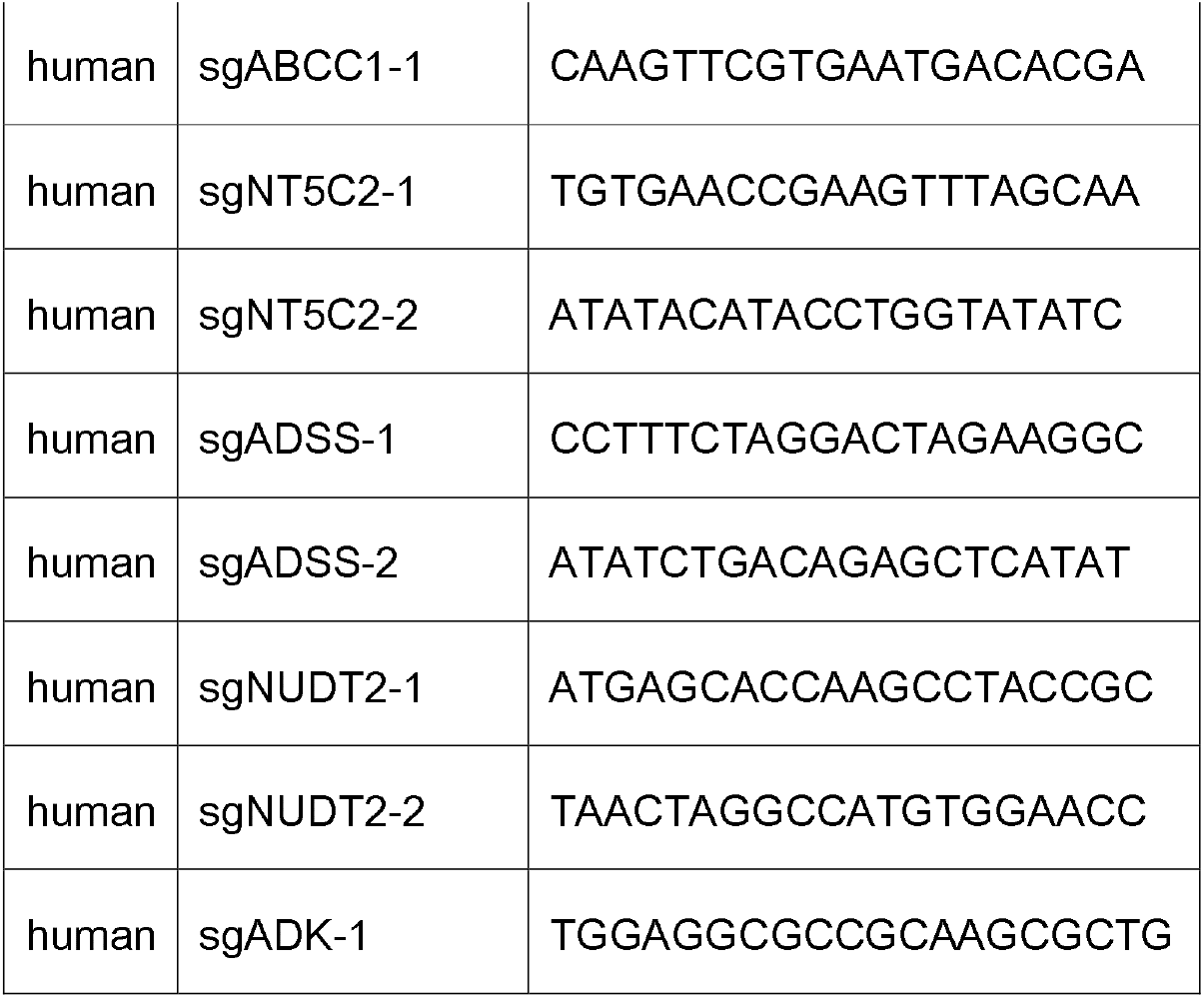
sequence of sgRNA used in validation experiments.

## Materials and Methods

### Cell lines and drugs

Eμ-Myc;Arf ^−/−^ cells were cultured in B-cell medium (45% DMEM, 45% IMDM and 10% FBS, supplemented with L-glutamate, and 5 μM β-mercaptoenthanol). Phoenix, HeLa, MDA-MB-231, A549, 293T and Huh7 were cultured in Dulbecco’s modified Eagle’s medium supplemented with glutamate and 10% (v/v) FBS. Remdesivir, ribavirin, favipiravir, chloroquine phosphate and hydroxychloroquine were provided by or purchased from MedChemExpress. Colchicine was purchased from Targetmol. Dexamethasone was purchased from Selleck.

### Plasmids and generation of stable cell lines

Murine or human NT5C2 cDNA and their activating mutant forms were synthesized by Genescripts. Cas9 was cloned to a retroviral vector. About 1% of Eμ-Myc;Arf ^−/−^ was infected by such Cas9-GFP retrovirus, from which we established a Cas9-expressing Eμ-Myc;Arf ^−/−^ stable cell line. Sequences of sgRNAs used in validation experiments are listed in Extended Data Table 3.

### Transfer of sgRNA libraries to a retroviral backbone

Eμ-Myc;Arf ^−/−^ cells cannot be efficiently infected with lentivirus. Therefore, we bulk transferred sgRNA sequences from the lentiviral library (Addgene: #1000000096) into a pBabe-SV40-puro backbone. Such a backbone can efficiently express sgRNAs and cause gene editing in Cas9-expressing Eμ-Myc;Arf ^−/−^ cells. Next-generation sequencing confirmed the quality of the subclone library, with around 98% of sgRNAs still present.

First, the U6-sgRNA segment from LentiGuide-Puro(Addgene #52963) was PCR amplified and transferred to pBabe-SV40-puro to generate a pBabe-U6sgRNA-SV40-puro vector with NheI and XhoI cloning sites.

Next, a mouse KO sgRNA pooled library (Addgene: #1000000096, 10sgRNA per gene) was amplified at 1000X fold coverage, and the sgRNAs were digested with NheI/XhoI from the library. The insert was gel purified twice and concentrated using the GeneJET Gel Extraction Kit(thermo scientific #K0692). pBabe-U6sgRNA-SV40-puro vector was digested with NheI/XhoI. After ligating the vector and sgRNA inserts, 20 tubes of 5-alpha Electrocompetent E. coli (Takara,#9027) were transformed using electroporation and the subcloned library was obtained by combining 4 maxipreps. The library complexity was confirmed by streaking diluted bacteria onto plates and counting colonies. The total number of colonies was > 100x the size of the library.

To generate a mCherry-expressing sgRNA vector used in Fig.5a, the plasmid pBabe-U6sgRNA-SV40-puro was digested to remove puro fragment and the mCherry fragment was cloned into the plasmid to create pBabe-U6sgRNA-SV40-mCherry.

Q5 Hot Start High-Fidelity DNA Polymerase (NEB #M0531L) was used for all cloning PCR reactions.

### Generation of sgRNA-expressing retrovirus and drug screen

Retrovirus packaging was achieved in phoenix cells using standard protocols. Around 100 million of phoenix cells were transfected with retroviral sgRNA library (10sgRNA per gene). Virus-containing medium was collected from these cells 2 days after transfection, and added to 230 million of Eμ-Myc;Arf ^−/−^ cells for infection. Infected cells gained puromycin resistance from the retroviral backbone, which enabled us to determine that around 23 million of Eμ-Myc;Arf ^−/−^ cells were infected. At such an infection rate, most cells should only contain one sgRNA. 23 million of infected cells also roughly correlate to 100X coverage of the sgRNA library.

After expansion of these infected cells, they were divided and 25 million of such cells were each subjected to treatment with a drug. Around 48hrs after treatment, cell viability dropped to 10~20%. Cells were centrifugated and resuspended in normal culture medium to allow for recovery. 3~5 days after this, cells reached viability between 60% to 70%. They were again treated with drugs and similar protocols was followed to remove drugs after this second round of treatment. After recovery from the second round of drug selection, 25 million of alive cells were collected. 25 million of no-treatment cells were also collected as a control.

Genomic DNA of the collected cells was extracted with Magen HiPure Tissue DNA Midi Kits (D3122-03), and sgRNA sequences were PCR amplified using the Q5® Hot Start High-Fidelity 2X master Mix. From each sample, 40μg of genomic DNA was used for PCR reaction. 8 tubes of 50μl PCR reaction was set up for each sample. PCR was stopped at 23 cycles. Water control was used to ensure there was no aerosol contamination.

The 400μl PCR products were purified with thermo scientific GeneJET Gel Extraction Kit (K0692) and dissolved in 30μl dH2O. Such products were separated with 2.5% agarose gel to obtain the 250bp PCR products. After purification with thermo scientific GeneJET Gel Extraction Kit (K0692), the composition of sgRNAs was analyzed by NGS at Novagene (Beijing). After NGS, Analysis of raw sequencing reads was performed using MAGeCKFlute analysi^*60*^. Fig. 2a and similar figures were generated from MAGeCKFlute analysis.

To generate the data for the type of analysis in 2A’, we performed the following analysis. If a gene knockout induced by sgRNA1.1 protected cells from a drug, this sgRNA1.1 will be enriched in the drug-selected population. Before drug selection, sgRNA1.1 made up 0.0025‰ of all sgRNAs from non-treated samples. After drug selection, that percentage increased to 1‰. From such data, we can conclude that sgRNA1.1 was enriched by 400 fold after drug selection. Similar method was used if a sgRNA was relatively depleted after drug selection.

### GFP or mCherry-based survival competition assay to assess relative resistance index conferred by cDNA expression or sgRNA knockout

Such experiments were used to produce data for Fig.4C’ and Fig.5A and 5E. The overall experimental design was previously described in Jiang *et al*^17^. Briefly, NT5C2 cDNA or its activating mutant along with GFP was stably expressed in a small portion of Eμ-Myc;Arf ^−/−^ cells. Remdesivir was applied to such cells at concentration that would kill 80~90% of uninfected parental cells. Since NT5C2 inactivates remdesivir and protected cells from remdesivir-induced cell death, after drug treatment, most of the surviving cells will be GFP positive that express NT5C2. If the GFP percentage rose from 20% in untreated cells to 80% in remdesivir-treated cells, it can be calculated that the **relative resistance index** is (0.8-0.8*0.2)/(0.2-0.8*0.2)=16, meaning that at such drug concentration, NT5C2-expressing cells are 16 fold as likely to survive remdesivir compared with control cells. The rationale behind such calculation and the biological meaning of relative resistance index is detailed in Jiang *et al* ^17^.

Certain events, such as ABCC1 knockout sensitizes cells to the killing by remdesivir. Therefore, sgABCC1 expressing, mCherry positive cells will die more after remdesivir treatment. In such scenario, we may observe that the mCherry positive rate drops from 50% in untreated cells to 20% in remdesivir-selected samples. From such numbers, it can be calculated that the **relative resistance index** is (0.2-0.2*0.5)/(0.5-0.5*0.2)=0.25, meaning ABCC1 knockout cells are **4 fold** as likely to die from remdesivir-induced killing. Accordingly, we will designate a **relative sensitization index** of **−4** to sgABCC1.

### Virus production

Zika virus stock was prepared as previously described^44^. Briefly, *in vitro* transcribed viral RNA was electroporated into Vero E6 cells, 72h-120h supernatants was harvested and titrated by plaque forming assay after removal of dead cells and cell debris.

### Infection of Huh7 derived cell line

Cells were seeded in 48-well plates for 24 hours before incubation with Zika virus at multiplicity of infection (MOI) equal to 1. 4 hours later, the unbound viruses were replaced by fresh medium containing remdesivir or ribavirin. After 24 hours, cells were collected and lysed by TRIZOL(Tiangen) to extract RNA. Such RNA samples were subjected to reverse transcription (Toyobo) and ZIKV genome copies were analyzed by quantitative PCR(Toyobo) with primers targeting ZIKV NS3. The sequences for the qPCR primers are GTTTGGCTGGCCTATCAGGT (forward) and GTGTCTGGTCCACACCTCTG (reverse).

